# Bioengineering a plant NLR immune receptor with a robust binding interface towards a conserved fungal pathogen effector

**DOI:** 10.1101/2024.01.20.576400

**Authors:** Rafał Zdrzałek, Yuxuan Xi, Thorsten Langner, Adam R. Bentham, Yohann Petit-Houdenot, Juan Carlos De la Concepcion, Adeline Harant, Motoki Shimizu, Vincent Were, Nicholas J. Talbot, Ryohei Terauchi, Sophien Kamoun, Mark J. Banfield

## Abstract

Bioengineering of plant immune receptors has emerged as a key strategy for generating novel disease resistance traits to counteract the expanding threat of plant pathogens to global food security. However, current approaches are limited by rapid evolution of plant pathogens in the field and may lack durability when deployed. Here, we show that the rice nucleotide-binding, leucine-rich repeat (NLR) immune receptor Pik-1 can be engineered to respond to a conserved family of effectors from the multihost blast fungus pathogen *Magnaporthe oryzae*. We switched the effector binding and response profile of the Pik NLR from its cognate rice blast effector AVR-Pik to the host-determining factor Pwl2 by installing a putative host target, OsHIPP43, in place of the native integrated HMA domain (generating Pikm-1^OsHIPP43^). This chimeric receptor also responded to other PWL alleles from diverse blast isolates. The crystal structure of the Pwl2/OsHIPP43 complex revealed a multifaceted, robust interface that cannot be easily disrupted by mutagenesis, and may therefore provide durable, broad resistance to blast isolates carrying PWL effectors in the field. Our findings highlight how the host targets of pathogen effectors can be used to bioengineer new recognition specificities that have more robust properties compared to naturally evolved disease resistance genes.

## Introduction

Engineering plant intracellular nucleotide-binding leucine rich repeat (NLR) immune receptors to generate new disease resistance profiles is an emerging method to expand the recognition capabilities of the plant immune system (1–3). NLRs orchestrate responses to pathogen virulence proteins (effectors) that are translocated into hosts during infection. Structure-guided bioengineering of the effector-binding regions of NLRs is a promising mechanism to modify receptor recognition specificity (4–8). Editing or domain-swapping of non-canonical integrated domains found in some NLRs has been particularly effective for either expanding or altering the effector response profiles of these immune receptors (4, 7, 9–13).

Engineering of NLR integrated domains for new effector specificities has been most extensively investigated in the rice paired NLRs RGA5/RGA4 and Pik-1/Pik-2. These NLR pairs confer resistance to blast fungus (*Magnaporthe oryzae*) strains carrying the AVR1-CO39/AVR-Pia or AVR-Pik effectors, respectively. These effectors are recognised through direct interaction with integrated Heavy Metal Associated (HMA) domains embedded in the sensor NLRs RGA5 or Pik-1 (14–17). Effector recognition by the NLR sensor results in receptor activation and initiation of defence responses that are dependent on their helper NLRs RGA4 and Pik-2 respectively (18–20). Intriguingly, each of these effectors are members of the sequence-divergent but structurally conserved family of *Magnaporthe* Avrs and ToxB-like (MAX) effectors (21). Computational structure prediction has shown the MAX family forms a large proportion of the *M. oryzae* effector repertoire (22) and the fold is overrepresented in effectors known to be detected by NLR immune receptors (14–16, 23).

The RGA5-HMA domain has been engineered to gain binding to non-corresponding MAX effectors including AVR-PikD and AVR-Pib (4, 9, 12, 13). Importantly, to achieve full resistance to AVR-Pib in cereals, the engineered full-length RGA5/RGA4 receptor pair required additional modification of the C-terminal region that directly follows the HMA domain in the RGA5 receptor (12, 13). Likewise, the HMA domain of Pik-1 can be engineered for new effector recognition specificity (5, 7, 9). The Pik NLRs and AVR-Pik effectors exist in allelic series, with different alleles of the receptors having different specificities towards effector variants (10, 15, 18, 24). Mutation of the Pik-1 HMA domain allows for expanded recognition of AVR-Pik effectors in Pik alleles with otherwise limited effector recognition spectrums (10, 24). Further, structure-guided studies demonstrate novel resistance can be generated against *M. oryzae* carrying stealthy variants of AVR-Pik by resurfacing the Pik-1 HMA to mimic that of the putative host target OsHIPP19 (7, 25). The Pik-1 HMA domain can also be exchanged for alternative HMA domains or other unique integrations (e.g. VHH nanobodies) to change the effector recognition specificity and modulate immune auto-activity (7, 9, 11).

Pathogen-encoded host-determinant factors are promising targets for generating novel resistance as they represent barriers to pathogen infection in certain host species that may be transferable (26). Pathogenicity towards Weeping Lovegrass 2 (Pwl2) is an effector from *M. oryzae* that is a host determinant factor for infection of weeping lovegrass (27, 28). Several variants of Pwl2 exist in *M. oryzae* populations. Two Pwl2 variants, Pwl2-2 and Pwl2-3 (which differ in only one or four amino acid positions of 145 in the full Pwl2 sequence), are not recognised by weeping lovegrass, overcoming the Pwl2-host determining barrier (28, 29). Further, Pwl2 is a member of a larger PWL effector family, including Pwl1, Pwl3 and Pwl4 that share 42-79% amino acid sequence identity with Pwl2 (30). Recently, natural resistance against Pwl2 has been identified in barley, conferred by the Mla3 (Rmo1) protein (27), but has not been identified in other cereal crop hosts of the blast fungus. Interestingly, Mla3 only recognises Pwl2 and not the Pwl2-2 variant. To date, the virulence function of Pwl2 (and other Pwl effector family members) during infection remains unclear, but as a host determinant of *M. oryzae* infection, it represents a promising target for receptor engineering.

In this study, we engineered the Pikm-1 sensor NLR by replacing the native HMA domain with the HMA-domain of the putative Pwl2 host target OsHIPP43 (31). We show that, in combination with Pik-2, this receptor responds to Pwl2 in *Nicotiana benthamiana*. Pwl2 binds OsHIPP43 with nanomolar affinity in vitro, and a crystal structure of the complex reveals an extensive interface formed between the proteins that proves challenging to disrupt by mutagenesis. The structure confirms Pwl2 as a MAX effector (27, 32), but with an additional C-terminal extension comprising an α-helix and loop region lacking secondary structure. Both the MAX fold and C-terminal extension are involved in OsHIPP43 binding. The engineered Pikm-1^OsHIPP43^/Pik-2 receptor also responds to Pwl2 allelic variants, and more divergent Pwl family members, raising the potential of this receptor to confer resistance to diverse *M. oryzae* strains carrying Pwl effectors, including the pandemic wheat blast lineage (33–37). Our study highlights the potential of using host-determinant factors as targets for new resistance strategies, and further demonstrates the strength of using host targets as effector recognition modules when integrated into plant NLRs.

## Results

### OsHIPP43 specifically binds Pwl2 in a yeast 2-hybrid library screen

We previously generated two libraries for yeast 2-hybrid (Y2H) screening, one comprising 195 *M. oryzae* effector candidates (38), and another comprising 151 putative Poaceae host target HMA domains (39, 40). Using these libraries, we conducted an all-vs-all screen in which we identified the rice HMA domain-containing protein Os01g0507700 (OsHIPP43) as a candidate interactor for Pwl1, Pwl2, and Pwl3 (38). To confirm the specificity of this interaction, we tested Pwl2 against a range of HMA-containing proteins from several HMA families that were either closely or distantly phylogenetically related (41) using pairwise Y2H assays (**Fig. 1A**). In this screen, Pwl2 interacted specifically with OsHIPP43, and no other HMA proteins (**Fig. 1A**). In contrast to Pwl2, we also reconfirmed AVR-PikD to be an effector with a promiscuous host HMA protein interaction profile (39, 41), including OsHIPP43 (**Fig. S1**).

**Fig. 1.**
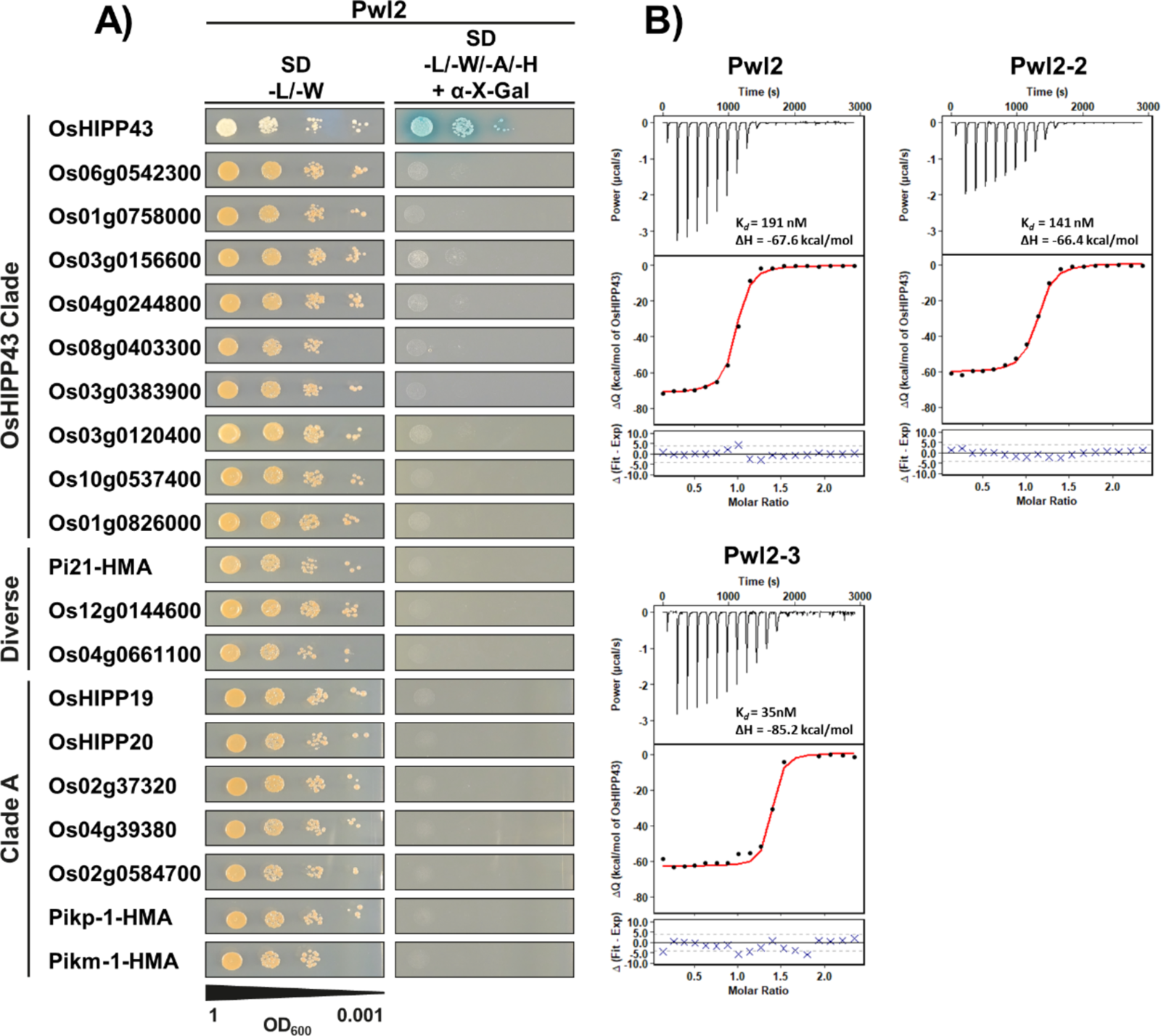
Pwl2, Pwl2-2 and Pwl2-3 bind OsHIPP43 with high affinity. **A)** Yeast 2-hybrid shows Pwl2 interacts specifically with OsHIPP43, but not with other tested HMA proteins across the HMA phylogeny. Blue colonies on selective medium (-L/-W/-A/-H + X-α-gal) indicate positive interactions. **B)** Binding affinity between Pwl2 and allelic variants with OsHIPP43 in vitro, as measured by ITC. **Top panels-** Representative raw isotherm showing heat exchange upon the series of injections of OsHIPP43 into the cell containing the effector. **Middle panels-** Integrated peaks from technical replicates and global fit to a single site binding model as calculated using AFFINImeter. **Bottom panels-** Difference between predicted value of measurement (by global fit) and actual measurement, as calculated using AFFINImeter.

### OsHIPP43 and Pwl2 effectors form high affinity complexes in vitro

To further characterise the interaction between OsHIPP43 and Pwl2, we performed isothermal titration calorimetry (ITC) to obtain binding affinities. To enable this, Pwl2 without the signal peptide (residues 22-145) and the HMA domain of OsHIPP43 (residues 26-101, henceforth referred to as OsHIPP43) were produced in *E. coli* and purified using a combination of immobilised-metal affinity chromatography (IMAC) and size-exclusion chromatography (SEC). OsHIPP43 interacted with Pwl2 with a calculated dissociation equilibrium constant (K*_d_*) of 191 nM, and high heat exchange upon binding, indicated by a change in enthalpy (ΔH) of -67.6 kcal/mol (**Fig. 1B**). To assess whether sequence polymorphisms in Pwl2 variants (**Fig. S2**) affect interaction with OsHIPP43 in vitro, we also expressed, purified and determined binding affinities for Pwl2-2 and Pwl2-3. We found Pwl2-2 and Pwl2-3 both bound OsHIPP43 with similar affinities to Pwl2, with K*_d_* values of 141 and 35 nM, respectively (**Fig. 1B**). These results confirm the initial interaction of OsHIPP43 and Pwl2 identified by Y2H analysis, and show that Pwl2 allelic variants bind OsHIPP43 with nanomolar affinity in vitro.

### Integration of OsHIPP43 into the Pikm-1 receptor switches recognition from AVR-Pik to Pwl2

Having established Pwl2 binds OsHIPP43 with high affinity, we hypothesised Pik-1 could be engineered to recognise Pwl2 in *N. benthamiana* by replacing the integrated HMA domain of the sensor Pikm-1 for the HMA domain of OsHIPP43.

To test this, we generated a Pikm-1^OsHIPP43^ chimera using Golden Gate cloning in a Pikm DOM2 ID-acceptor vector (9). Co-expression of Pikm-1^OsHIPP43^ with the Pikm-2 helper NLR in *N. benthamiana* resulted in an effector-independent cell death, indicative of auto-activity (**Fig. S3**). To circumvent this, we tested a sensor/helper allelic mismatching strategy of Pik-2 helpers, as described in (9, 42), by co-expressing Pikm-1^OsHIPP43^ with the Pikp-2 helper. By contrast to co-expression with Pikm-2, co-expression of Pikm-1^OsHIPP43^ with the Pikp-2 helper did not result in effector-independent cell death (**Fig. S3**).

Next, we tested whether the chimeric Pikm-1^OsHIPP43^/Pikp-2 receptor could respond to Pwl2 when co-expressed in *N. benthamiana*. Co-expression of Pikm-1^OsHIPP43^/Pikp-2 with Pwl2 resulted in a robust cell death response, indicating the chimeric receptor is able to recognise Pwl2 (**Fig. 2A**, **Fig. S4**). This cell death response was dependent on co-expression with Pikp-2. Importantly, no cell death was observed on co-expression with AVR-PikD demonstrating a switch in sensor NLR specificity.

**Fig. 2.**
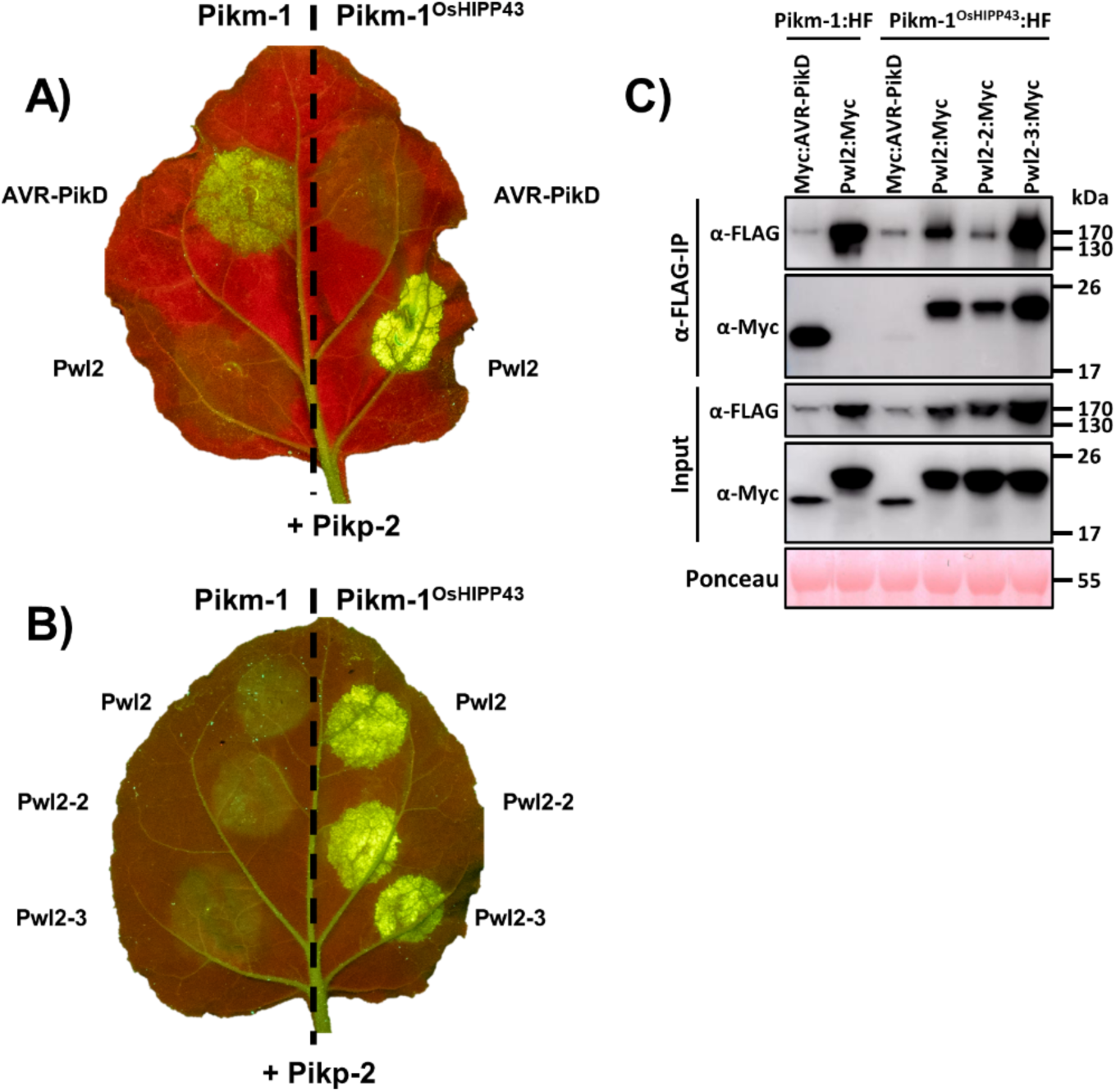
The Pikm-1^OsHIPP43^/Pikp-2 chimera recognises Pwl2 allelic variants on expression in *N. benthamiana*, underpinned by direct binding. **A, B)** Cell death assays showing Pwl2, Pwl2-2 and Pwl2-3, but not AVR-PikD, are recognised by the chimeric Pikm-1^OsHIPP43^/Pikp-2 receptor. Leaves were imaged under UV light, allowing visualisation of cell death responses as green fluorescence. **C)** Co-immunoprecipitation assay showing chimeric Pikm-1^OsHIPP43^ receptor association with Pwl2, Pwl2-2 and Pwl2-3. All proteins were transiently expressed in *N. benthamiana* via agroinfiltration. **Upper panel-** Anti-FLAG immunoprecipitation (αFLAG-IP) was followed by Western Blot detection with relevant antibodies. **Lower panel-** Input confirms presence of all proteins prior to immunoprecipitation. Ponceau staining was used to demonstrate even protein loading.

As Pwl2 allelic variants Pwl2-2 and Pwl2-3 interacted with OsHIPP43 in vitro, we tested whether Pikm-1^OsHIPP43^/Pikp-2 could respond to these effectors in planta. Co-expression of Pikm-1^OsHIPP43^/Pikp-2 with either Pwl2-2 or Pwl2-3 resulted in a robust cell death response in both cases, equivalent to Pwl2 (**Fig. 2B**, **Fig. S4**). Taken together, these data show that the chimeric Pikm-1^OsHIPP43^/Pikp-2 receptor recognises three Pwl2 allelic variants, generating an NLR immune receptor with novel recognition specificity.

### The chimeric Pikm-1^OsHIPP43^ receptor associates with Pwl2 and allelic variants in planta

To determine whether the *N. benthamiana* cell death responses are likely underpinned by direct protein interactions between the effectors and the NLRs, we performed a co-immunoprecipitation (co-IP) assay. For this, we transiently co-expressed either Pikm-1 or Pikm-1^OsHIPP43^ with epitope tags (HellFire; 6xHis, 3xFLAG) alongside Myc-tagged effectors. Subsequently, we performed anti-FLAG pulldowns and probed for the presence of the differentially tagged proteins. As previously demonstrated, wild-type Pikm-1 associated with AVR-PikD, but did not associate with Pwl2 (**Fig. 2C**). Correlating with the cell death assay results, Pwl2, Pwl2-2 and Pwl2-3 associated with Pikm-1^OsHIPP43^ in planta (**Fig. 2C**).

Unexpectedly, we also observed association of AVR-PikD with the Pikm-1^OsHIPP43^ chimera (**Fig. 2C**), which does not translate to a cell death response, and AVR-PikD/OsHIPP43 binding was not observed in vitro (**Fig. S5A, B**). However, as mentioned above, we did observe an interaction between AVR-PikD and OsHIPP43 by Y2H. We also tested the interaction of AVR-PikD with Pikm-1 in the absence of an HMA domain (Pikm-1^ΔHMA^), and still observed association, suggesting a certain level of association is not dependent on binding to the HMA domain (**Fig. S5C, D**). Similarly, HMA-independent binding of AVR-Pia to an RGA5 mutant with its HMA domain removed was previously observed, yet this binding did not trigger a downstream immune response (4). Together, these co-IP data demonstrate that recognition of Pwl2 alleles by the Pikm-1^OsHIPP43^ chimera is underpinned by protein-protein interactions in planta, and the sensor NLR’s HMA domain is required for immune signalling.

### Pwl variants bind OsHIPP43 and are recognised by the chimeric Pikm-1^OsHIPP43^/Pikp-2 receptor in planta

Pwl1, Pwl3 and Pwl4 share 42-79% percent amino acid sequence identity with Pwl2 (excluding the signal peptide) (**Fig. S6**) and would therefore be expected to adopt a similar protein structure. We hypothesised these effectors might also interact with OsHIPP43 in vitro. To test this, we expressed and purified Pwl1 and Pwl4 from *E. coli* and used ITC to measure their binding affinity to OsHIPP43. Pwl1 and Pwl4 interacted with OsHIPP43 with similar affinities as Pwl2, with K*_d_* values of 147 nM and 124 nM, respectively (**Fig. 3C**). We were not able to produce Pwl3 in sufficient quantities for in vitro binding experiments, so it was excluded from this analysis.

**Fig. 3.**
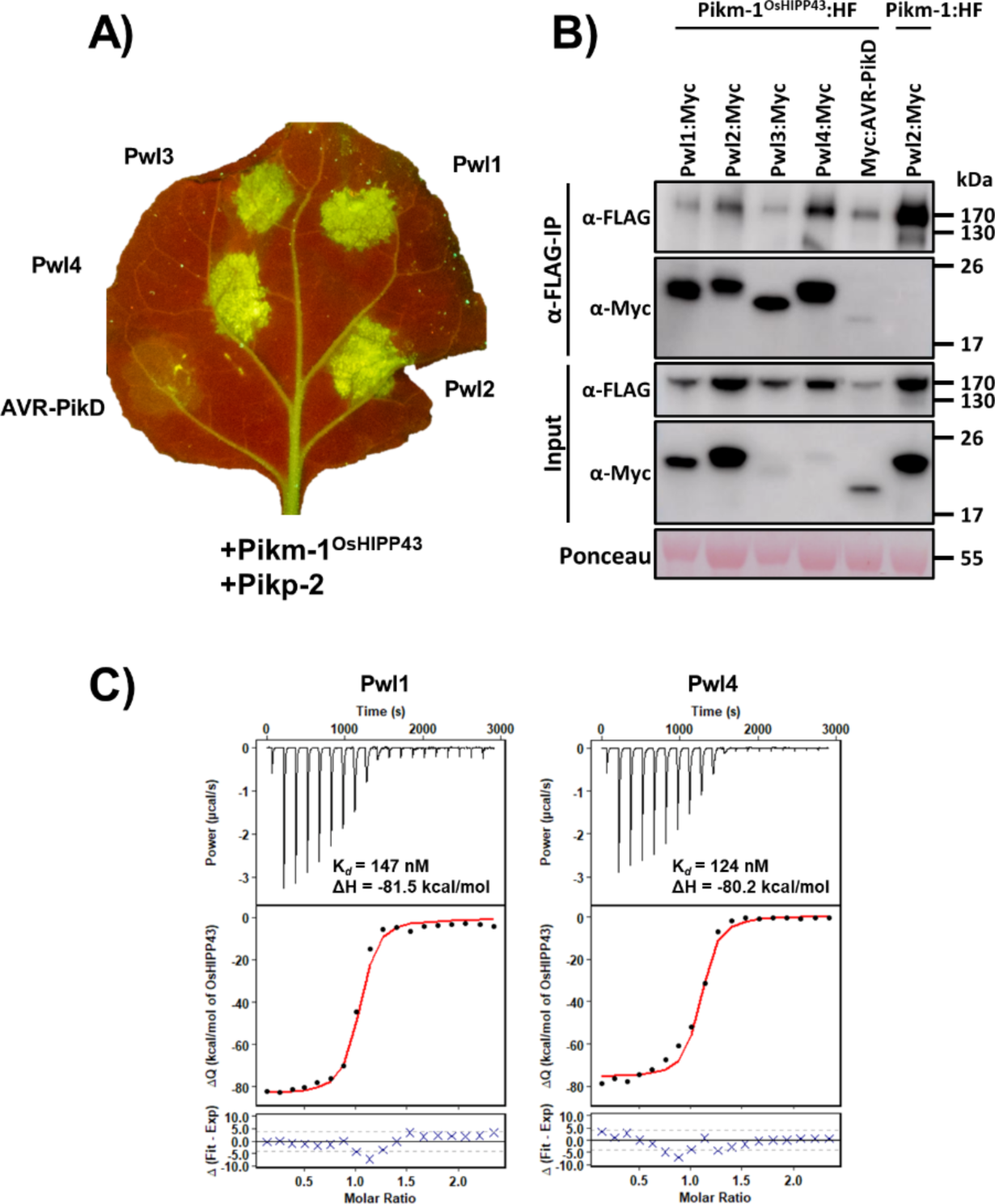
The Pikm-1^OsHIPP43^/Pikp-2 chimera recognises Pwl effector variants on expression in *N. benthamiana*, underpinned by direct binding. **A)** Cell death assay showing Pwl variant recognition by the chimeric Pikm-1^OsHIPP43^/Pikp-2 receptor. Leaves were imaged under UV light, allowing visualisation of cell death responses as green fluorescence. **B)** Co-immunoprecipitation assay showing chimeric Pikm-1^OsHIPP43^ receptor association with Pwl effectors. All proteins were transiently expressed in *N. benthamiana* via agroinfiltration. **Upper panel-** Anti-FLAG immunoprecipitation (αFLAG-IP) was followed by Western Blot detection with relevant antibodies. **Lower panel-** Input confirms presence of all proteins prior to immunoprecipitation. Ponceau staining was used to demonstrate even protein loading. **C)** Binding affinity between Pwl variants and OsHIPP43 in vitro as measured by ITC. **Top panels-** Representative raw isotherm showing heat exchange upon the series of injections of the OsHIPP43 into the cell containing the effector. **Middle panels-** Integrated peaks from technical replicates and global fit to a single site binding model as calculated using AFFINImeter. **Bottom panels-** Difference between predicted value of measurement (by global fit) and actual measurement, as calculated using AFFINImeter.

To test whether direct OsHIPP43 binding observed for Pwl1 and Pwl4 can result in recognition in planta, we transiently co-expressed these effectors (and Pwl3) in the presence of Pikm-1^OsHIPP43^/Pikp-2 in *N. benthamiana*. We observed a strong cell death response for all Pwl effector variants tested (**Fig. 3A, Fig. S7**), indicating recognition by the Pikm-1^OsHIPP43^ receptor. To test whether Pwl effectors associate with Pikm-1^OsHIPP43^ in planta, we performed co-IP experiments as described previously. We found all Pwl effectors associated with the chimeric Pikm-1^OsHIPP43^, despite weak accumulation of some effectors in planta (**Fig. 3C**). Taken together, in vitro binding of the Pwl effector variants to OsHIPP43 correlates with in planta co-IP and cell death assays, indicating the Pikm-1^OsHIPP43^ receptor can directly interact with the wider family of Pwl effectors to mediate recognition.

### Pwl2 adopts a MAX effector fold structure and forms an extensive interface with OsHIPP43

To understand the structural basis of interaction between Pwl2 and OsHIPP43, we determined the crystal structure of the effector/target complex by *X*-ray crystallography. For crystallisation, a stable form of the Pwl2/OsHIPP43 complex was identified by limited proteolysis with trypsin (**Fig. S8**). Mass spectrometry of the digested sample revealed a ten amino acid truncation at the C-terminus of Pwl2 (Pwl2^Δ10^), while OsHIPP43 remained intact. We then cloned this truncation of Pwl2, co-expressed in *E. coli* with OsHIPP43, and purified the complex. After sparse matrix screening, protein crystals were obtained in 1.2 M potassium sodium tartrate tetrahydrate, 0.1 M Tris pH 8.0. *X*-ray diffraction data were collected from these crystals at the Diamond Light Source, UK, to a resolution of 1.8 Å. We also collected a highly redundant long wavelength *X*-ray dataset at λ=1.9 Å to enable structure solution using a sulphur-single anomalous dispersion (S-SAD) approach.

In the structure of the Pwl2^Δ10^/OsHIPP43 complex (henceforth the Pwl2/OsHIPP43 complex), the HMA domain of OsHIPP43 adopts the well-characterised HMA fold, comprising two α-helices and a four-stranded antiparallel β-sheet. Interestingly, the two cysteines, Cys-39 and Cys-42, form a disulphide bridge within the conserved metal binding motif (MDCEGC) that faces away from the interaction interface with Pwl2 (**Fig. 4A, B**). For Pwl2, the structure reveals three predominant features. Firstly, an N-terminal region (residues Trp-25 to Pro-85) adopts the MAX fold, a conformation repeatedly observed for experimentally determined structures of *M. oryzae* effectors (15–17, 23, 43, 44). Unusually for structurally characterised MAX effectors, this region is followed by an α-helix (residues His-87 to His-100) and finally a C-terminal region devoid of major secondary structure features (**Fig. 4A, B**).

**Fig. 4.**
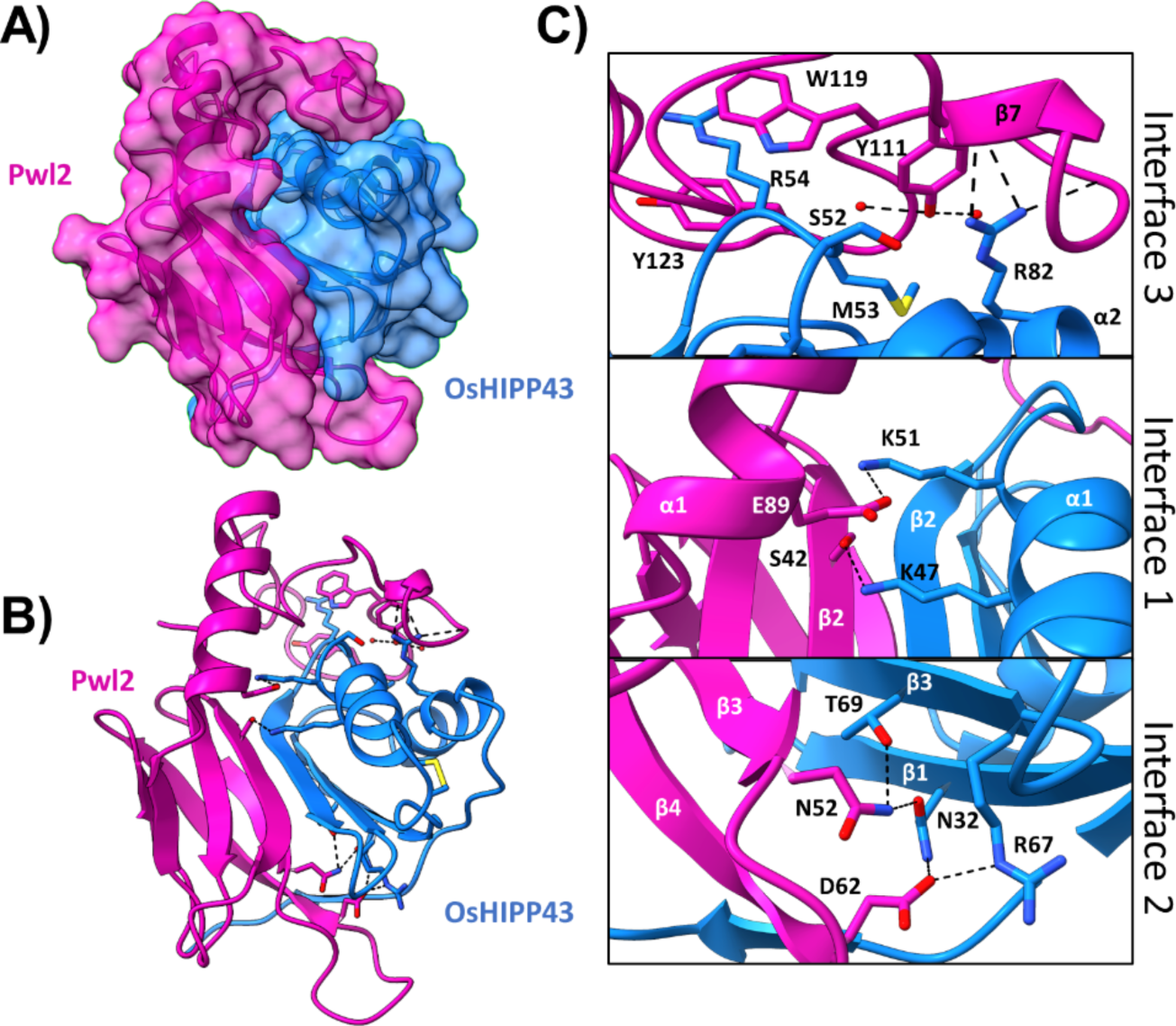
Crystal structure of the Pwl2/OsHIPP43 complex reveals an extensive interface. **A)** Transparent surface representation of Pwl2 (pink) and OsHIPP43 (blue), with secondary structures displayed. **B)** With surfaces hidden, Pwl2 can be seen wrapping around OsHIPP43, forming an extensive interface. Key residues for protein-protein interaction are shown in stick representation. **C)** Close-up views of molecular interactions across three interfaces, as described in the text, with side chains shown in stick representation. α-helices and Δ-strands are labelled, and amino acids are labelled with single letter codes. Hydrogen bonds are depicted as black dashes between atoms. Red spheres represent water molecules.

Pwl2 and OsHIPP43 interact via an extensive interface (**Fig. 4A, B**). Analysis with QtPisa (45, 46) revealed that 25.1% and 38.2% of accessible protein surface area is buried in the complex for Pwl2 and OsHIPP43, respectively, with a total interface area of 1976.9 Å^2^. For comparison, the total interaction interface formed by the complex of Pikp-1-HMA/AVR-PikD is 966.6 Å^2^ (15). More than half of the residues in Pwl2 and OsHIPP43 (62/112 and 45/76, respectively) contribute to the interaction, which can be divided into three distinct interfaces (**Fig. 4B, C**). Interface 1 includes Pwl2 residues Ser-42 and Glu-89 that form hydrogen bonds with Lys-47 and Lys-51, respectively, located on the α1 helix of OsHIPP43. Interface 2 is predominantly formed by the loop between β3 and β4 of Pwl2. Residues Asn-52 and Asp-62 of Pwl2 form hydrogen bonds with OsHIPP43 residues Asn-32, and Arg-67 and Thr-69 located on β3 of OsHIPP43 (**Fig. 4B, C**). Finally, interface 3 is formed by the C-terminal region of Pwl2 that folds across the structure of OsHIPP43. In this region, Pwl2:Tyr-111 interacts with OsHIPP43 residues Ser-52 and Met-53, positioned on a loop between α1 and β2, and Arg-82, located on α2. Further, Trp-119 and Tyr-123 form π-stacking interactions with the hydrophobic chain of OsHIPP43:Arg-54 (**Fig. 4B, C**). The structure of the Pwl2/OsHIPP43 complex is a novel example of how MAX effectors bind HMA proteins, which continue to emerge as a major host targets of *M. oryzae* effectors (**Fig. S9**).

### Recognition of Pwl2 by Pikm-1^OsHIPP43^ is robust to single point mutation

To validate the structure and explore the limits of the chimeric Pikm-1^OsHIPP43^/Pikp-2 receptor to recognise Pwl2, we performed site-directed mutagenesis followed by in planta cell death assays in *N. benthamiana*. With the aim of disrupting complex formation, we designed seven individual point mutants of Pwl2, dispersed across the three Pwl2/OsHIPP43 interfaces. These included S42R and E89R at interface 1, N52R and D62R at interface 2, and Y111R, Y119D and W123D at interface 3 (single amino acid codes are used to describe mutants). Surprisingly, a robust cell death response was retained on co-expression of Pikm-1^OsHIPP43^/Pikp-2 with each of the seven individual Pwl2 mutants (**Fig. 5A**, **Fig. S10**). We concluded that single point mutations are not sufficient to overcome Pwl2 recognition by the Pikm-1^OsHIPP43^/Pikp-2 receptor.

**Fig. 5.**
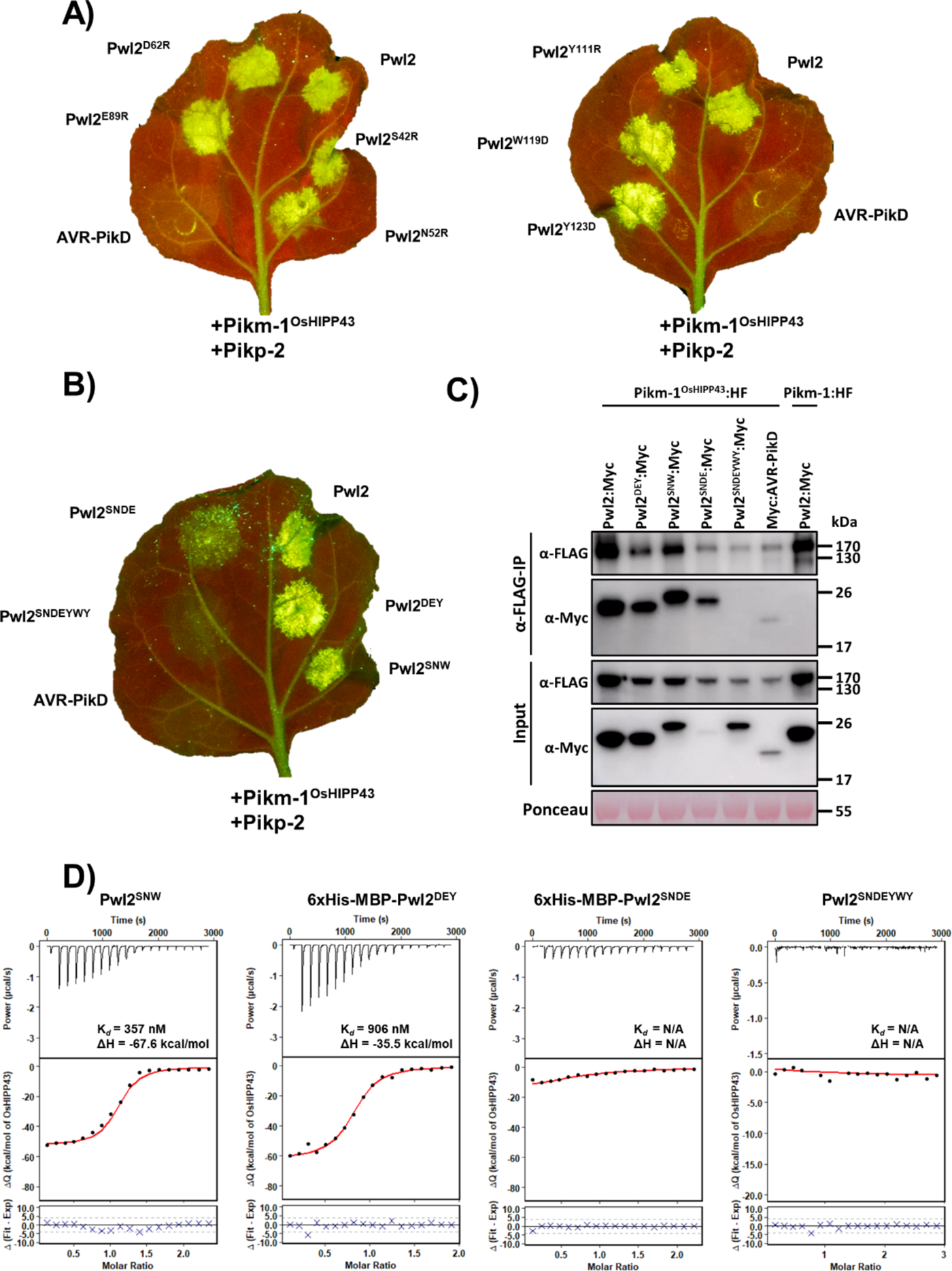
The Pwl2/Pikm-1^OsHIPP43^/Pikp-2 response and Pwl2/Pikm-1^OsHIPP43^ and Pwl2/OsHIPP43 interaction are robust to single or multiple point mutations. **A)** Cell death assays showing recognition of all single Pwl2 point mutants by the chimeric Pikm-1^OsHIPP43^/Pikp-2 receptor, despite deliberate targeting of mutations at the Pwl2/OsHIPP43 interface. **B)** Cell death assays showing recognition of multiple structure-led mutants of Pwl2 by the chimeric Pikm-1^OsHIPP43^/Pikp-2 receptor, as described in the text. Leaves were imaged under UV light, allowing visualisation of cell death responses as green fluorescence. **C)** Co-immunoprecipitation assay showing chimeric Pikm-1^OsHIPP43^ receptor association with all tested Pwl2 mutants, except for the septuple Pwl2^SNDEYWY^ mutant. **Upper panel-** Anti-FLAG immunoprecipitation (αFLAG-IP) was followed by Western Blot detection with relevant antibodies. **Lower panel-** Input confirms presence of all proteins prior to immunoprecipitation. Ponceau staining was used to demonstrate even protein loading. **D)** Binding affinity between multiple Pwl2 mutants and OsHIPP43 in vitro, as measured by ITC. **Top panels-** Representative raw isotherm showing heat exchange upon the series of injections of the OsHIPP43 into the cell containing the effector. **Middle panels-** Integrated peaks from the technical replicates and global fit to a single site binding model as calculated using AFFINImeter. **Bottom panels-** Difference between predicted value of measurement (by global fit) and actual measurement as calculated using AFFINImeter. Amino acids are labelled with single letter codes throughout.

### Combined mutations across Pwl2/OsHIPP43 interfaces are not sufficient to abolish recognition and binding

Next, we designed two triple mutants in Pwl2, targeting one residue at each of the previously identified interfaces, in an attempt to disrupt recognition and binding by the Pikm-1^OsHIPP43^ receptor. These mutations combined either D62R, E89R and Y123D (Pwl2^DEY^), or S42R, N52R and W119D (Pwl2^SNW^). Both Pwl2^DEY^ and Pwl2^SNW^ mutants were still recognised by the chimeric Pikm-1^OsHIPP43^/Pikp-2 receptor in *N. benthamiana* cell death assays (**Fig. 5B**, **Fig. S11**), and retained interaction with Pikm-1^OsHIPP43^ when tested by co-IP (**Fig. 5C**). To investigate whether these triple mutants affected the strength of binding, we expressed and purified the proteins from *E. coli* and tested for interaction with OsHIPP43 in vitro using ITC. Pwl2^SNW^ was purified as for the wild-type effector, but Pwl2^DEY^ could only be obtained in sufficient quantities for analysis without cleaving the 6xHis-MBP (Maltose Binding Protein) tag. We confirmed that the presence of 6xHis-MBP did not affect the ITC assay by measuring the affinity of tagged wild-type Pwl2 (6xHis-MBP-Pwl2) with OsHIPP43, showing it was comparable to untagged Pwl2, and that 6xHis-MBP alone did not bind OsHIPP43 (**Fig. S12**). We determined the binding affinities of 6xHis-MBP-Pwl2^DEY^ and Pwl2^SNW^ with OsHIPP43 as K*_d_* = 906 nM and K*_d_* = 357 nM, respectively (**Fig. 5D**). While the affinity of Pwl2^SNW^ with OsHIPP43 is broadly comparable to wild type, Pwl2^DEY^ displays reduced affinity, possibly due to the partial instability of the protein. Nonetheless, both mutants show tight binding to OsHIPP43 in vitro. We therefore concluded that these triple mutants, despite involving multiple regions in the Pwl2/OsHIPP43 interface, are not sufficient to break the complex with OsHIPP43.

### Evasion of recognition requires extensive disruption of the Pwl2/OsHIPP43 interface

Finally, we combined four mutations: S42R, E89R (interface 1) and N52R and D62R (interface 2) to generate a quadruple mutant (Pwl2^SNDE^). We also generated a septuple mutant combining all mutations described above into one protein (Pwl2^SNDEYWY^). When transiently co-expressed in planta, the Pwl2^SNDEYWY^ mutant was not recognised by the Pikm-1^OsHIPP43^/Pikp-2 receptor (**Fig. 5B, Fig. S11**), and did not associate with Pikm-1^OsHIPP43^ in co-IP experiments (**Fig. 5C**), despite accumulating to detectable levels. Pwl2^SNDEYWY^ was stable when expressed and purified from *E. coli*. We tested for interaction of this mutant with OsHIPP43 using ITC. Consistent with co-IP data, we did not observe any interaction (**Fig. 5D**). Intriguingly, co-expression of the Pwl2^SNDE^ mutant with Pikm-1^OsHIPP43^ displayed reduced cell death in *N. benthamiana* compared to wild-type Pwl2 (**Fig. 5B**, **Fig. S11**). This potentially results from reduced accumulation of Pwl2^SNDE^ in planta, as association of this mutant with Pikm-1^OsHIPP43^ was still observed in a co-IP assay (**Fig. 5C**). We purified limited amounts of Pwl2^SNDE^ with an uncleaved 6xHis-MBP tag, but we could not detect binding between this mutant and OsHIPP43 by ITC, possibly due to overall low stability of the protein. (**Fig. 5D**). However, co-expression of Pwl2^SNDE^ with OsHIPP43 enabled purification of a complex that was crystallised. These crystals diffracted *X*-rays generating a dataset to 2.8 Å resolution. In the resulting structure, we find that the S42R, N52R, D62R and E89R mutations are all accommodated via a rearranged protein interface. At interface 1, the introduced Arg-42 and Arg-89 are rotated relative to the smaller Ser and Glu residues present in the wild-type protein and positioned away from OsHIPP43 residues Lys-51 and Lys-47 with which they previously formed hydrogen bonds. Interestingly, this leads to formation of an alternative hydrogen bond between Lys-47 of OsHIPP43 and Ser-40 from Pwl2 (**Fig. S13A**). At interface 2, the introduction of Arg-52 and Arg-62 disrupts the position of the loop from Tyr-53 and Arg-63 (**Fig. S13B**) with the consequent removal of hydrogen bonds and hydrophobic interactions observed in the wild-type complex. At interface 3, the Pwl2^SNDE^/OsHIPP43 and Pwl2/OsHIPP43 complexes adopt a very similar structure.

Overall, based on the results of mutagenesis, we conclude the interaction between Pwl2 and OsHIPP43 is not easily compromised, even using structure-guided approach. Indeed, partial and full loss of OsHIPP43 binding and recognition in planta requires multiple mutations across the extensive interface.

## Discussion

Engineering plant NLR immune receptors is a promising strategy to develop novel resistance to plant diseases (47). As our knowledge of NLR engineering expands, the integration of host targets as sensor domains for effectors has emerged as a powerful tool for creating new effector recognition specificities (3). In this study, we engineered recognition of the non-host resistance factor Pwl2 from the blast pathogen *M. oryzae*. By incorporating a host target of Pwl2, OsHIPP43, into the Pikm-1 receptor chassis, we generated a receptor that binds and responds to Pwl2 allelic variants and related Pwl effectors in planta. Further, we define Pwl2 as a MAX effector, and demonstrate it binds the OsHIPP43 HMA domain through a novel extended MAX effector fold, which uses a binding interface previously unobserved in other MAX effector/HMA complexes. Taken together, this study highlights the potential of host targets as effector recognition modules that can be incorporated into NLRs, mimicking the evolution of naturally occurring integrated domains, and is a promising approach to generate novel resistance to disease.

*M. oryzae* strains expressing Pwl2 are unable to infect weeping lovegrass and this effector therefore acts as a host-specificity barrier (28, 30). This non-host resistance trait allowed for Pwl2 cloning nearly 30 years ago (28), but only recently has an immune receptor capable of recognising Pwl2 been characterised in barley, the NLR Mla3 (Rmo1) (27). However, biophysical and structural studies of Pwl2 (and its allelic variants/relatives) have been limited, largely due to absence of a defined or putative virulence target, and the protein presenting as disordered in solution (48). We hypothesised that knowledge of host target interactions for Pwl2 could inform bioengineering efforts seeking to exploit NLR integrated domains as baits for effectors (4, 7, 9, 11–13), and provide opportunities for expanded resistance profiles. For example, Mla3 (Rmo1) does not provide resistance to *M. oryzae* strains expressing Pwl2-2 (27).

The role of OsHIPP43 as a putative virulence-associated target of Pwl2 has been explored in a companion paper (29). Here, we focussed on understanding how the interaction between Pwl2 and OsHIPP43 could be used for immune receptor engineering. The crystal structure of Pwl2 bound to OsHIPP43 reveals Pwl2 to be a MAX effector, with a C-terminal helical extension and extended loop region. Each of these regions contribute to an extensive interface with OsHIPP43, but at least the C-terminal region is likely to be disordered in solution in the absence of a binding partner. This is consistent with previous biophysical analyses of Pwl2 (48), and may explain why structural studies of this protein have not been described to date. Previously, the interfaces between MAX effectors (such as AVR-Pik and AVR-Pia/AVR1-CO39) and integrated HMA domains (including those from the NLRs Pik-1 and RGA5) were shown to be spatially distinct (15, 17, 49), implying multiple interaction modes that can support immune recognition via HMA domains. The Pwl2/OsHIPP43 interaction defines a third mode that is distinct, but spatially similar, to that observed in the AVR1-CO39/RGA5-HMA complex (**Fig. S9A, B**), also incorporating the novel C-terminal extension of Pwl2.

Interestingly, the polymorphic residues found in Pwl2 allelic variants Pwl2-2 and Pwl2-3 (at positions Glu-89, Asp-90, Lys-91 and Ser-92) are not all directly located at the interface with OsHIPP43 (**Fig. S14**). These residues do not impact binding to OsHIPP43 in vitro or in planta, and each Pwl2 variant is recognised by the Pikm-1^OsHIPP43^/Pikp-2 receptor in cell death assays. Unlike Pwl2, Pwl2-2 and Pwl2-3 do not define a barrier for *M. oryzae* infection of weeping lovegrass, suggesting that this non-host resistance is not based on HIPP43 interaction (for example, via interaction with the weeping lovegrass OsHIPP43 homologue).

Studies involving the Pik-1 receptor as a chassis for novel integrated domains have successfully altered effector recognition specificity. The AVR-Pik host target OsHIPP19 can be incorporated into Pikp-1 to expand the recognition specificity of the receptor to stealthy variants of AVR-Pik, including in transgenic rice (7). Recently, VHH nanobodies, which share no sequence or structural homology with HMA domains, could be integrated into Pikm-1 to generate bespoke ligand specificity to non-effector targets (9, 11). The work presented here further demonstrates the utility of the Pik-1 chassis to incorporate new domains for recognition of diverse pathogen effectors.

The Pikm-1^OsHIPP43^/Pikp-2 receptor is not only capable of recognising Pwl2, Pwl2-2 and Pwl2-3, but also the other Pwl variants, Pwl1, Pwl3 and Pwl4, that are present in diverse, host-adapted lineages of *M. oryzae*. Further, it proved to be difficult to break the interaction between Pwl2 and OsHIPP43 through structure-led mutagenesis. Only mutagenesis across at least two interfaces was capable of impairing recognition. This suggests the deployment of the Pikm-1^OsHIPP43^/Pikp-2 receptor in host plants may confer disease resistance challenging to overcome through point mutation. Indeed, overcoming of Pikm-1^OsHIPP43^-mediated resistance may require Pwl effectors to be deleted, which could potentially impact pathogen fitness and virulence (29). As exemplified by Flor’s gene-for-gene model (50), arms-race co-evolution of host/pathogen interactions posits that selection pressure on pathogens can drive the evolution of effectors with mutations to avoid detection by the host immune system. For example, natural variants of the blast (*M. oryzae*) AVR-Pik effector and stem rust (*Puccinia graminis f. sp. tritici*) AvrSr50 effector escape NLR recognition through substitution of surface exposed residues (10, 25, 51). However, effector mutation could be detrimental to putative virulence functions if it also results in loss of host target interactions. Using host targets as integrated domains to underpin recognition has the advantage that effector mutation to avoid detection may also result in loss of binding to host targets and reduce virulence.

As discussed above, the engineered Pikm-1^OsHIPP43^/Pikp-2 immune receptor not only responds to Pwl2 and close allelic variants, but also to other PWL family members. Surveys of available *M. oryzae* genomes from strains that infect a wide range of cereal hosts, including the wheat blast pandemic lineage that is an emerging threat global wheat production (52, 53), demonstrate the almost universal presence of these Pwl effectors in the pathogen population. While future work is required to confirm that Pikm-1^OsHIPP43^/Pikp-2 can confer broad disease resistance against diverse *M. oryzae* strains carrying Pwl effectors in cereal hosts, this work highlights the potential of host-target-led immune receptor engineering for agriculture.

## Materials and Methods

### Cloning for in-planta expression

Wild type Pikm-1, Pikm-2, Pikp-2 and AVR-PikD were cloned as described in (10). Pikm-1^ΔHMA^ was cloned using the Golden Gate system, by fusing the CC domain directly to the NB-LRR domains, mas promoter and terminator in the pICH47751 acceptor vector. The chimeric construct Pikm-1^OsHIPP43^ used for co-IP experiments, was cloned in the DOM2 acceptor vector, as described in (9). Briefly, Pikm-1 was assembled in a LVL0 acceptor vector, with the sequence of the HMA domain exchanged for an RFP selection cassette, flanked by BpiI restriction sites. This allowed for exchange of the selection cassette for the sequence of OsHIPP43 and subsequent cloning into a LVL1 construct, under control of the mas promoter. For cell death assays, we generated a Golden Gate LVL2 acceptor, including a hygromycin selection cassette, 2xCaMV35S:Pikm-1^RFP^:HF (as described above, but with BsaI flanking sites), and mas:Pikp-2:HA in the pICSL4723 backbone. Next, we used Golden Gate cloning to introduce the OsHIPP43 or wild-type Pikm-1 HMA domain into the Pikm-1 sequence. Pwl effectors were cloned with a C-terminal 4xMyc tag into pICH47751, under control of the AtUbi10 promoter. All Pwl2 mutants were commercially synthesised by Integrated DNA Technologies as a gBlocks gene fragments.

### Cloning for recombinant expression in *E. coli*

Pwl and OsHIPP43 sequences were cloned into the pOPIN-GG vector pPGN-C (54) with a cleavable N-terminal 6xHis, 6xHis-GB1 or 6xHis-MBP tag using the Golden Gate system. For co-expression and crystallisation with Pwl2 variants 6xHis-MBP-Pwl2, 6xHis-Pwl2^Δ10^ or 6xHis-GB1-Pwl2^SNDE^, OsHIPP43 was cloned into pPGC-K (54) without a tag using the Golden Gate system.

### Y2H

We utilized the Matchmaker Gold Yeast Two-Hybrid System (Takara Bio USA) to investigate the interactions between HMA domain-containing proteins and Pwl2. The DNA sequences encoding HMA domain-containing proteins were inserted into the pGBKT7 vector and co-transformed with Pwl2 (in the pGADT7 vector), into chemically competent *Saccharomyces cerevisiae* Y2HGold cells (Takara Bio USA). Following transformation, single colonies grown on selection plates were inoculated into 5 ml of SD-Leu-Trp medium and incubated overnight at 28 °C to an optical density at 600 nm (OD_600_) of 1-1.5. The culture was then used to prepare 1:10 serial dilutions, starting from OD_600_ = 1. Subsequently, 5 μl of each dilution was spotted onto both an SD-Leu-Trp plate as a growth control and an SD-Leu-Trp-Ade-His plate containing X-α-gal, as described in the user manual. Following incubation at 28 °C for 72 hours, the plates were imaged. Each experiment was conducted a minimum of three times, yielding consistent outcomes.

### Protein expression and purification

Pwl2 and Pwl2-2 were purified with 6xHis affinity tags. All the other Pwl effectors (including Pwl2 mutants) and AVR-PikD were purified with 6xHis-GB1 tags (unless stated otherwise). Expression vectors were transformed into BL21-AI One Shot (Arabinose Inducible) *E. coli* cells (Invitrogen). The expression vector encoding 6xHis-MBP-OsHIPP43 was transformed into *E. coli* SHuffle cells (55). For inoculation, 5 ml of overnight pre-culture was added to 1 L of LB medium in 2 L baffled Erlenmeyer flask with appropriate antibiotics, which was then was incubated with shaking at 37 °C (BL21-AI strain) or 30 °C (SHuffle strain) until the OD reached 0.6 - 0.8. Then the temperature was decreased to 18 °C and cultures were induced with 0.2% arabinose (BL21-AI strain) or 1 mM IPTG (SHuffle strain) and incubated overnight (for at least 18 hours). 8 L of cultures were grown per construct.

For expression of Pwl2/OsHIPP43 and Pwl2^SNDE^/OsHIPP43 complexes, plasmids encoding the two proteins were transformed into the *E.* coli SHuffle strain as tagged (6xHis-MBP for Pwl2, 6xHis for Pwl2^1′10^, and 6xHis-GB1 for Pwl2^SNDE^) and untagged (OsHIPP43) proteins. Following expression, the complexes were purified as described below for individual proteins.

Cells were harvested by centrifugation (10 min, 7500 x g, 4 °C) and resuspended in ice-cold lysis buffer (50 mM HEPES pH 8.0, 500 mM NaCl, 5% glycerol, 50 mM glycine and 20 mM imidazole), freshly supplemented with cOmplete EDTA-free Protease Inhibitor Cocktail. Subsequently, the cells were lysed by sonication and the lysate was clarified by centrifugation at (25 min, 45000 x g, 4 °C).

The resulting supernatant was loaded onto ÄKTAxpress to perform immobilised metal affinity chromatography (IMAC), directly followed by size-exclusion chromatography (SEC) in SEC buffer (20 mM HEPES pH 7.5, 150 mM NaCl). Samples were then incubated overnight with recombinant 3C protease at 4 °C to remove the affinity/solubility tags. After cleavage, the untagged protein was separated from cleaved tags and the 6xHis-tagged 3C protease by affinity chromatography using 5 ml HisTrap HP NTA column (GE Healthcare). If proteins were also tagged with MBP, an MBPTrap HP dextrin sepharose column (GE Healthcare) was also used in tandem with the HisTrap column. The sample was further purified by SEC. Final samples were flash frozen in liquid nitrogen and stored at - 70 °C.

To purify proteins without cleaving the 6xHis-MBP tag, after the IMAC and SEC, samples were passed through 3xMBPTrap HP dextrin sepharose column (GE Healthcare) and subsequently eluted using SEC buffer supplemented with 10mM maltose. To purify 6xHis-MBP protein, the cleaved tag from the 6xHis-MBP-Pwl2 purification was removed from the solution using a MBPTrap HP dextrin sepharose column (GE Healthcare) and subsequently eluted using SEC buffer supplemented with 10mM maltose.

### Isothermal titration calorimetry (ITC)

ITC experiments were conducted using a MicroCal PEAQ-ITC (Malvern, UK). All protein samples were exchanged into the same buffer prior to each experiment via overnight dialysis in Slide-A-Lyzer MINI Dialysis Devices (Thermo Scientific). The Pwl effectors were placed in the experimental cell at 20 μM and titrated with OsHIPP43 at 200-300 μM at 25 °C. In each run, a single injection of 0.5 μL of OsHIPP43 was followed by 19 injections of 2 μl at 150 s intervals, with stirring at 750 rpm. Experiments were done in triplicate. The raw titration data was analysed using AFFINIMeter software (56) that integrated the datasets, removed noise, corrected the baseline and calculated the ΔH and Ka parameters. The K*_a_* parameter was then converted into K*_d_* using the operation K*_d_* =1/K*_a_*.

### Analytical size-exclusion chromatography

Analytical SEC experiments were carried out at 4 °C using a Superdex 75 10/300 GL column (GE Healthcare) equilibrated in SEC running buffer. AVR-PikD and OsHIPP43 were mixed in a 1:1 molar ratio and incubated on ice for 1 h. A sample volume of 110 μl was injected on the column. For analysis of individual proteins, samples (110 μl) were loaded at a concentration of 1 mg/ml. The samples passed through the column at a flow rate of 0.5 ml/min. 0.5 ml fractions were collected for SDS-PAGE analysis. The protein elution profile was monitored by measuring the absorbance at 280 nm.

### Tryptic digest

Stock solution of trypsin (Sigma) was prepared at 1 mg/ml in 1 mM HCl. A of 1:3 dilution series in PBS (137 mM NaCl, 2.7 mM KCl, 10 mM Na_2_HPO_4_, 1.8 mM KH_2_PO_4_, pH 7.4) was prepared using 5 μl of diluted trypsin solution per tube. To each tube, 20 μl of protein was added at concentration of 0.2 mg/ml. The samples were incubated at room temperature for 30 min. The reaction was stopped by adding 7 μl of stopping buffer (4x Loading Dye supplemented with 100 μM DTT and 5x cOmplete EDTA-free Protease Inhibitor Cocktail), and incubating at 95 °C for 10 min. 15 μl of the mix was loaded on a gel for the SDS-PAGE analysis.

### Crystallisation, *X*-ray data collection and structure solution of the Pwl2/OsHIPP43 complex

Crystallization trials were set up using the sitting drop, vapor diffusion method. Trials were set up in 96 well plates using an Oryx nano robot (Douglas Instruments, United Kingdom). Plates were kept at 20 °C. Crystals were obtained for the Pwl2^Δ10^/OsHIPP43 complex in the ProPlex^TM^ crystallisation screen (Molecular Dimensions) from 1.2 M Potassium sodium tartrate tetrahydrate, 0.1 M Tris 8.0, after ∼2 weeks at a protein concentration of 35 mg/ml. For data collection, crystals were harvested, cryoprotected in buffer containing the crystallisation condition supplemented with 20% ethylene glycol, and flash-frozen in liquid nitrogen.

*X*-ray diffraction data were collected at beamline I04 of the Diamond Light Source (Oxford, UK) under beamline proposal mx25108. Data reduction was carried out using the xia2.dials (Native data, (57)) and xia2.multiplex (S-SAD data, (58, 59)) pipelines with the scaled (but unmerged) data imported and processed with AIMLESS (as implemented in CCP4i2) (60, 61). The structure was solved by the Single-wavelength Anomalous Dispersion (SAD) method using the CRANK2 pipeline as implemented in CCP4i2 (62, 63). The resulting model was then used as a template to solve the high-resolution Native dataset by molecular replacement with PHASER (64). To arrive at the final structure, a series of manual rebuilding, refinement and validation steps were carried out using REFMAC (65) and COOT (66). The structure was validated with MolProbity (67) and tools implemented in COOT. Protein interface analysis was carried out using QtPISA (68). Final data collection, refinement, and validation statistics are shown in **Table S1** and **Table S2**. The final structure and the *X*-ray diffraction data used to derive this have been deposited at the Protein Data Bank with the accession number 8R7A. The structure was visualised for presentation using ChimeraX (69).

### Crystallisation, *X*-ray data collection and structure solution of Pwl2^SNDE^/OsHIPP43 complex

The crystallisation trials were set up as described above, at a concentration of 35 mg/ml. Crystals were obtained in the PEGSuite^TM^ screen from the condition 0.2 M ammonium formate, 20% PEG 3350. *X*-ray diffraction data was collected at beamline I04 at the Diamond Light Source (Oxford, UK) under beamline proposal mx25108. Data reduction was carried out using the xia2.dials (57) pipeline with the scaled (but unmerged) data imported and processed with AIMLESS (as implemented in CCP4i2) (60, 61). The structure was solved by the molecular replacement with PHASER (64) using the Pwl2/OsHIPP43 complex as a template. The final structure was generated and validated as described above for the Pwl2/OsHIPP43 complex. Final data collection, refinement, and validation statistics are shown in **Table S1** and **Table S2**. The final structure and the *X*-ray diffraction data used to derive this have been deposited at the Protein Data Bank with the accession number 8R7D. The structure was visualised for presentation using ChimeraX (69).

### Agroinfiltration

Prior to infiltration, *Agrobacterium tumefaciens* strain GV3101 (C58 (rifR) Ti pMP90 (pTiC58DT-DNA) (gentR) Nopaline (pSouptetR)) were transformed with constructs of interest by electroporation and grown for two days on LB plates with relevant antibiotics at 28 °C. Bacteria were gently scraped from the plate and resuspended in infiltration buffer (10 mM 2-(N-morpholine)-ethanesulfonic acid (MES) pH 5.6, 10 mM MgCl2, with freshly added 150 μM acetosyringone). Bacteria were mixed in desired combinations to give an OD_600_ in the final inoculum as following: agrobacteria carrying NLRs were infiltrated at OD_600_ = 0.4 and bacteria carrying the effectors at OD_600_ = 0.6. Each inoculum also contained agrobacteria carrying the p19 construct at OD_600_ = 0.1. The prepared agrobacteria were infiltrated into the leaves of 4-week-old *N. benthamiana* leaves using needleless 1 ml syringes.

### Cell death assays

Cell death assays were conducted as described in (20). Briefly, at 5 days post infiltration (dpi), detached agroinfiltrated *N. benthamiana* leaves were imaged under UV light (abaxial side). Each infiltration spot was scored (0 to 6) for cell death occurrence, according to the scale published in (15). Scoring data was plotted individually for each sample as dot plots using R v4.0.5. (https://www.r-project.org/) and the graphic package ggplot2 (70). Every dot represents an individual data point / score. All dots are randomly scattered around their given cell death score (with the size of the circle at a given score being proportional to the number of dots within). Each dot has a distinct colour corresponding to the biological replicate.

### Co-immunoprecipitation (co-IP) assays

Proteins of interest were transiently co-expressed in 4-to 5-week-old *N. benthamiana* plants following agroinfiltration, using OD_600_ = 0.4 for NLRs, OD_600_ = 0.6 for effectors, OD_600_ = 0.1 for p19. As an exception, bacteria carrying AVR-PikD were infiltrated at OD_600_ = 0.06, to even the level of expression between the effectors (strong expression of AVR-PikD otherwise dominated the Western Blots). At 3 dpi, five leaf discs were harvested into Eppendorf tubes and flash frozen in liquid nitrogen. Frozen leaf samples were ground into powder and resuspended in 650 µl cold extraction buffer (GTEN buffer (25 mM Tris-HCl (pH 7.5), 10% glycerol, 1 mM EDTA and 150 mM NaCl), supplemented with 0.1% NP-40 (Sigma), 0.5% w/v PVPP, 1x protease inhibitor cocktail (Sigma), 10 mM DTT and Roche protease inhibitor (50 ml/1 tablet)). Supernatants were collected by centrifugation (30 mins, 4 °C). 20 μl of each supernatant was collected then mixed with 20 μl of 2x laemmli buffer and denatured at 95 °C for 10 minutes (input samples). The remaining supernatant samples were mixed with 8 µl M2 anti-FLAG magnetic beads (Sigma, M8823) and incubated for 1 h on a rotary mixer. To remove non-specifically bound proteins, anti-FLAG magnetic beads were washed 5 times with cold wash buffer (GTEN buffer plus 0.1 % NP-40 (Sigma) and Roche protease inhibitor (50 ml/ 1 tablet)). For elution of proteins from the anti-FLAG beads, samples were incubated at 70 °C for 10 min with 40 μl 2x laemmli buffer (IP samples). Finally, all the input and IP samples were separated by SDS-PAGE.

### Western blot

Proteins from SDS-PAGE gels were transferred onto a PVDF (polyvinylidene difluoride) membrane (pre-activated in methanol for 1 min) using Trans-Blot Turbo transfer system (Bio-Rad) according to the manufacturer’s protocol. After protein transfer, the membrane was incubated with blocking buffer (5% w/v skimmed milk in TBS-T (50 mM Tris-HCl pH 8.0, 150 mM NaCl, 0.1% Tween-20)) at 4 °C for 1 h, with agitation. Subsequently, membranes were incubated with appropriate antibodies diluted in blocking buffer (α-FLAG: Cohesion Biosciences, at 1:3000 dilution; α-Myc: Santa Cruz Biotechnology, at 1:5000 dilution) overnight. The next day, membranes were washed with TBS-T and visualised using the LumiBlue ECL Extreme reagents (Expedeon) or Clarity Max Western ECL Substrate (Bio Rad) in the ImageQuant LAS 500 spectrophotometer (GE Healthcare). To visualise total protein loaded, membranes were stained with Ponceau red stain.

## Acknowledgements

We thank current and past members of the “BLASTOFF” team from the Banfield, Kamoun, Terauchi, Talbot and Moscou Laboratories for discussion and Emma Turley for reading of the manuscript. We also thank David Lawson, Clare Stevenson, and Julia Mundy for help with crystallisation and *X*-ray data collection. For funding, we thank the UKRI Biotechnology and Biological Sciences Research Council (BBSRC, UK, grants BB/W00108X/1, BB/WW002221/1, BB/V002937/1, BB/V016342, BB/V015508/1, BB/P012574, BBS/E/J/000PR9795, BBS/E/J/000PR9796 and BB/X010996/1), UKRI-BBSRC Norwich Research Park Biosciences Doctoral Training Partnership (grant BB/M011216/1), the European Research Council (743165 “BLASTOFF”, 101077853 “PANDEMIC”), the John Innes Foundation, the Gatsby Charitable Foundation, and the Japanese Society for the Promotion of Science (JSPS, KAKENHI 20H05681, 23K20042).

---

**Fig. S1.**
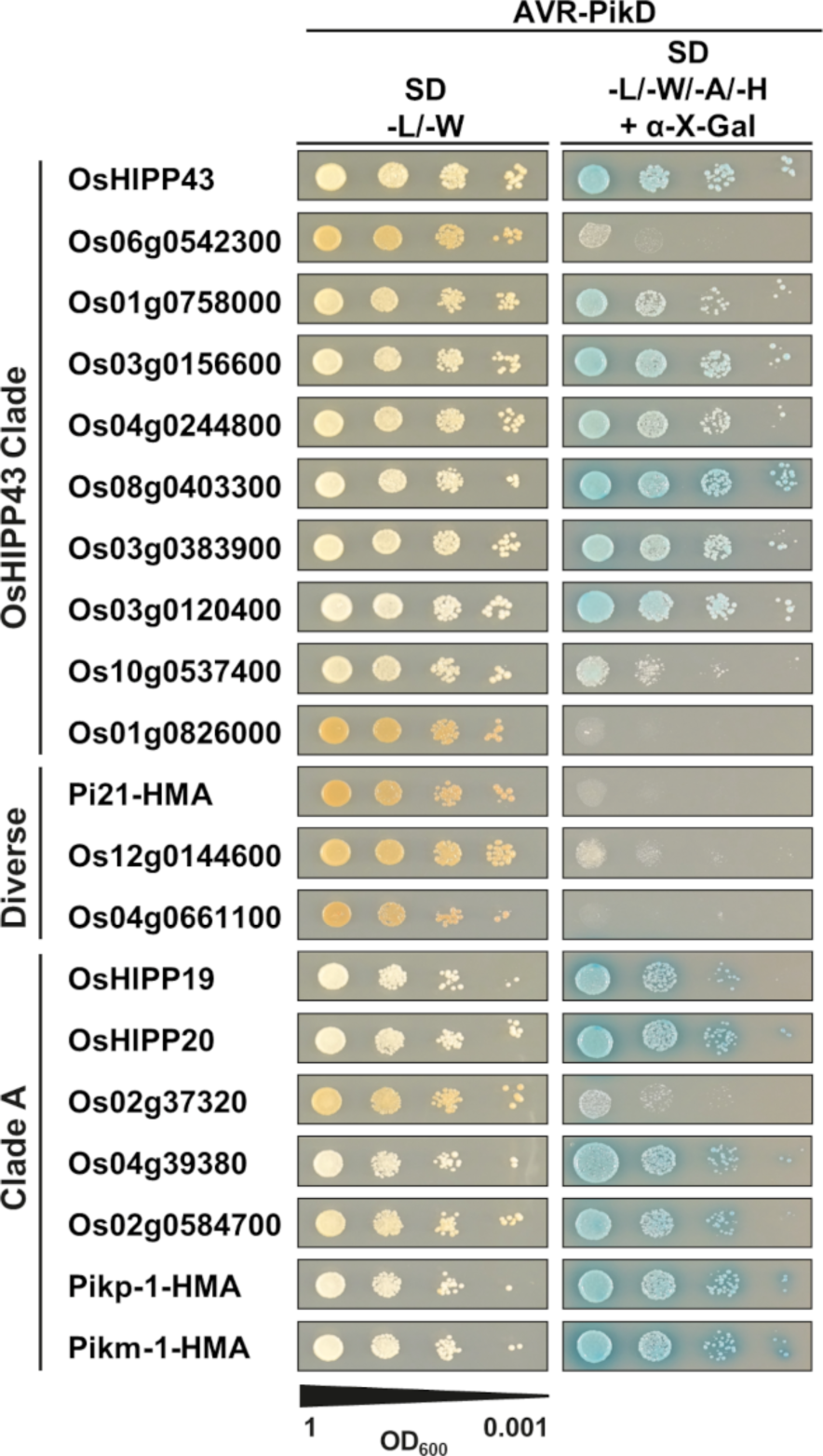
AVR-PikD interacts with different HMA domains in Yeast 2-hybrid screen. Yeast 2-hybrid shows AVR-PikD interacts with various HMA proteins across the HMA phylogeny. Blue colonies on selective medium (-L/-W/-A/-H + X-α-gal) indicate positive interaction.

**Fig. S2.**
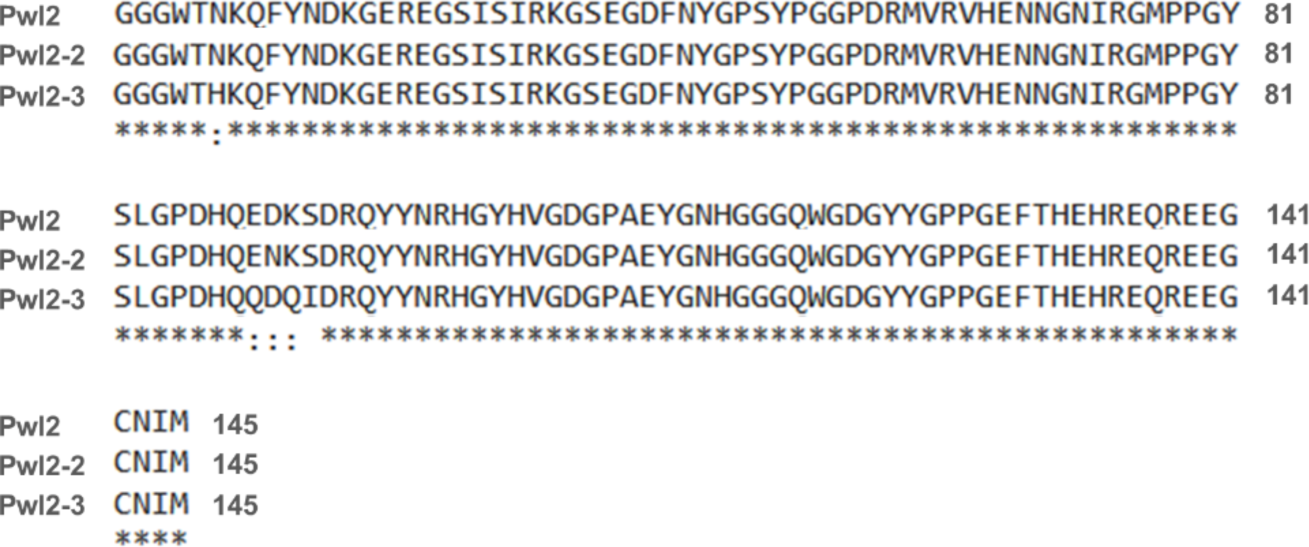
Sequence alignment of Pwl2 allelic variants identifies five polymorphic residues, with four clustered together. Alignment performed using Clustal Omega (https://www.ebi.ac.uk/Tools/msa/clustalo/). Signal peptides were removed from the alignment.

**Fig. S3.**
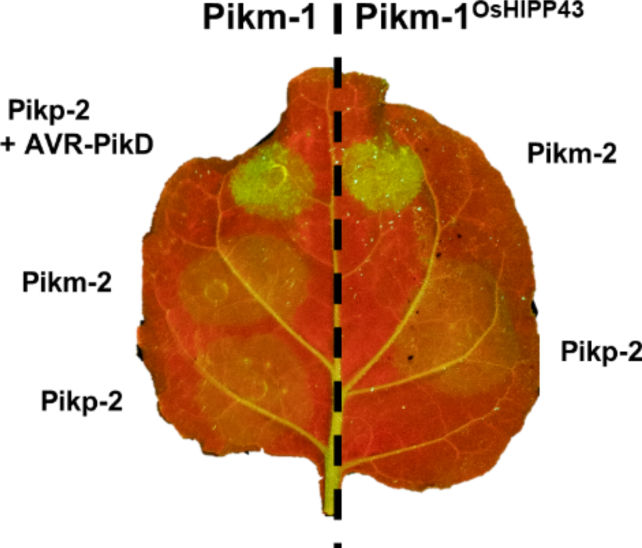
Auto-activity of Pikm-1^OsHIPP43^ with Pikm-2 can be alleviated by mismatching with Pikp-2. Cell death assays showing the chimeric Pikm-1^OsHIPP43^ receptor is auto-active when co-expressed with Pikm-2 in *N. benthamiana*, but not Pikp-2. Leaves were imaged under UV light, allowing visualisation of cell death responses as green fluorescence.

**Fig. S4.**
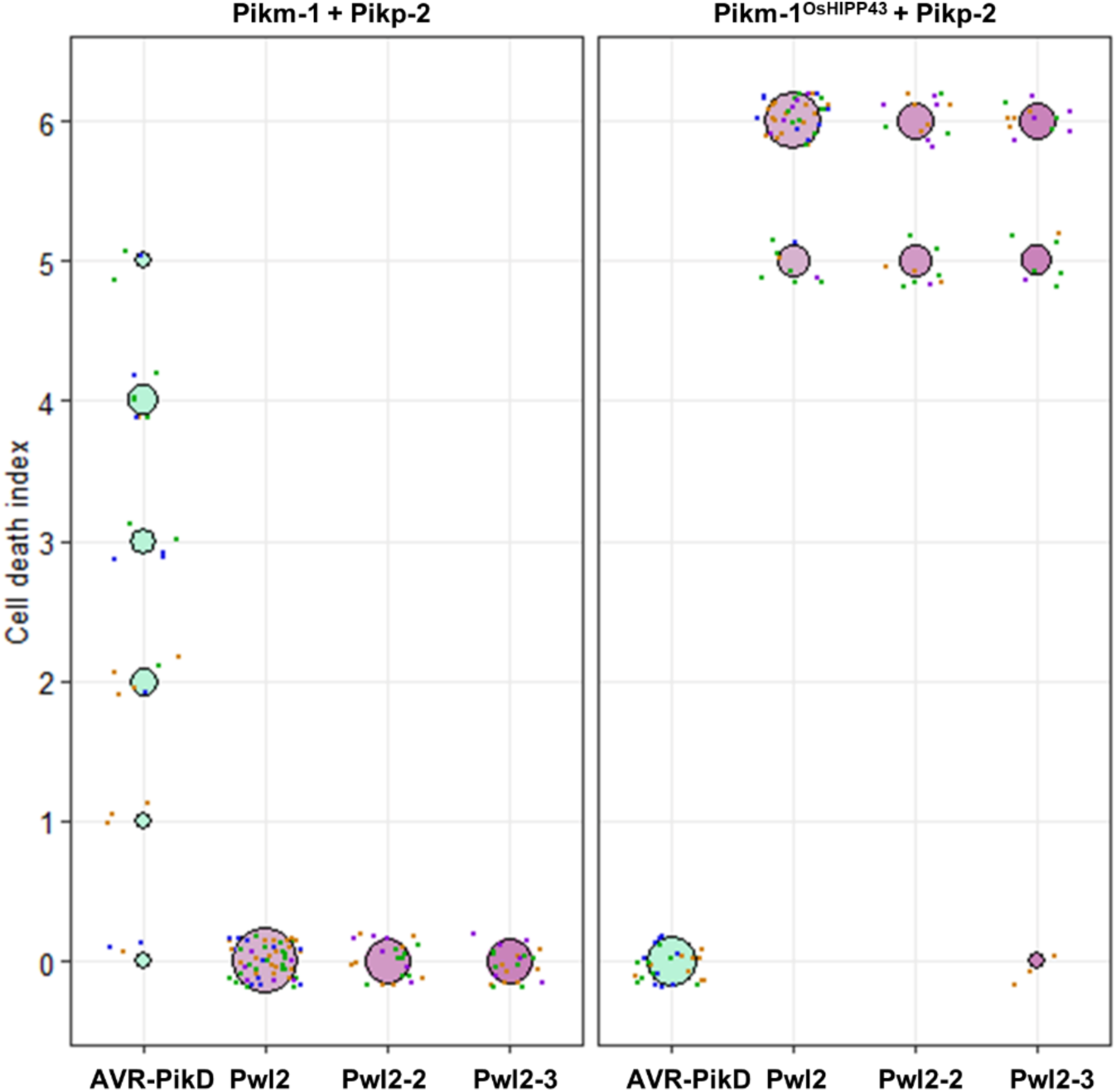
Quantification of cell death assays presented in Fig. 2. Each dot represents a single scoring point in the assay. All dots are randomly scattered around the cell death score for visualisation (size of the circle at given score is proportional to the number of dots within). The colour of each dot reflects independent biological replicates.

**Fig. S5.**
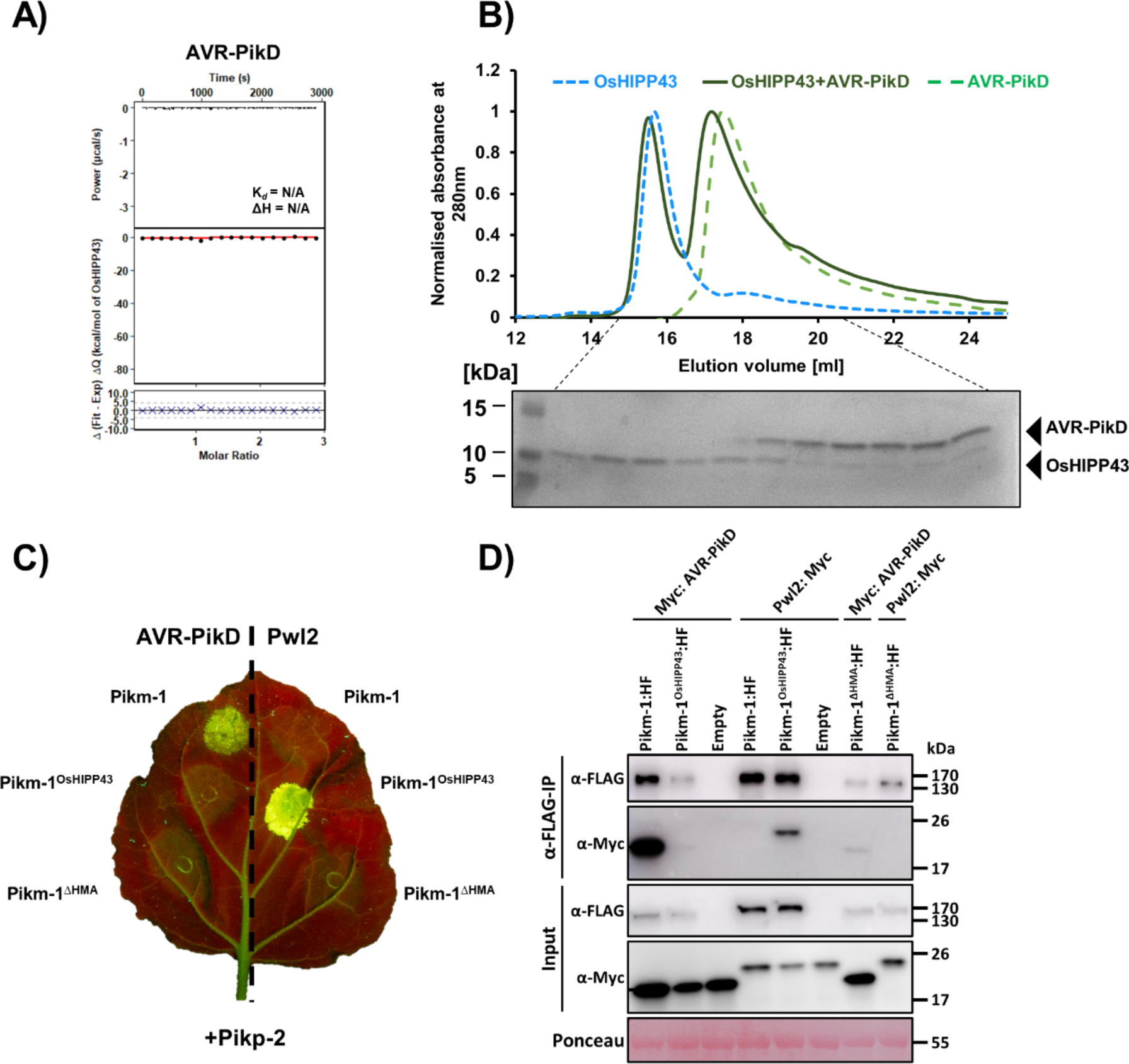
AVR-PikD does not interact with OsHIPP43 in vitro and is not recognised by the Pikm-1^OsHIPP43^/Pikp-2 receptor, but can interact with Pikm-1^OsHIPP43^ in planta. A) Titration of AVR-PikD with OsHIPP43 in ITC experiments shows a lack of binding between the proteins. **Top panel-** Representative raw isotherm showing heat exchange upon the series of injections of OsHIPP43 into the cell containing the effector. **Middle panel-** Integrated peaks from technical replicates and global fit to a single site binding model as calculated using AFFINImeter. **Bottom panel-** Difference between predicted value of measurement (by global fit) and actual measurement, as calculated using AFFINImeter. **B)** Following pre-incubation, AVR-PikD and OsHIPP43 elute as distinct peaks (equivalent to individual proteins) on analytical size-exclusion chromatography, supporting the lack of binding observed by ITC. **C)** AVR-PikD is not recognised by the chimeric Pikm-1^OsHIPP43^/Pikp-2 receptor, nor by the Pikm-1^ΔHMA^ receptor in the cell death assay. Leaves were imaged under UV light, allowing visualisation of the cell death responses as green fluorescence. **D)** Pwl2 specifically interacts with the chimeric Pikm-1^OsHIPP43^ NLR in co-IP experiments, and AVR-PikD can non-specifically interact with chimeric Pikm-1 receptors, independent of the HMA region. All proteins were transiently expressed in *N. benthamiana* via agroinfiltration. **Upper panel-** Anti-FLAG immunoprecipitation (αFLAG-IP) was followed by Western Blot detection with relevant antibodies. **Lower panel-** Input confirms presence of all proteins prior to immunoprecipitation. Ponceau staining was used to demonstrate even protein loading.

**Fig. S6.**
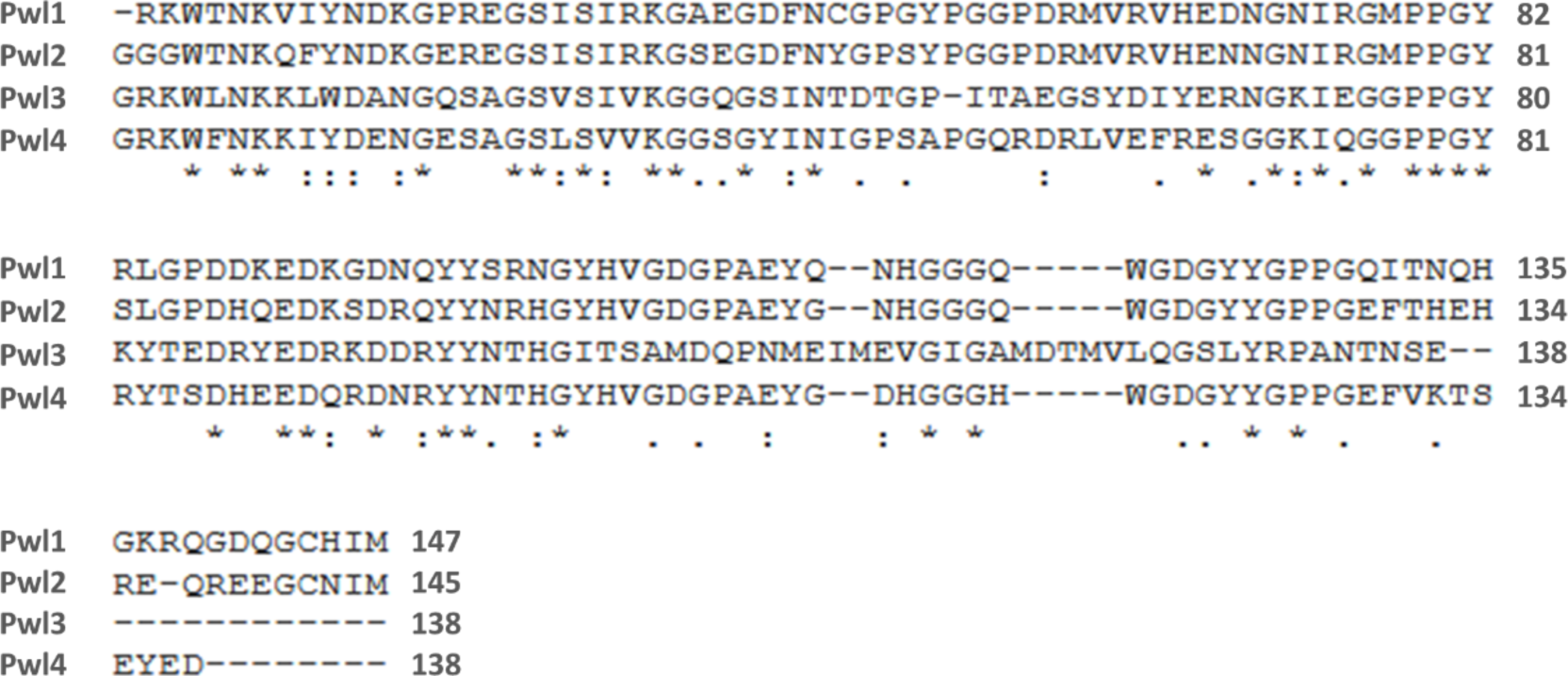
Pwl variants share between 41 and 79% sequence identity. Sequence alignment of the Pwl family effectors, revealing extensive sequence diversity. Alignment performed using Clustal Omega (https://www.ebi.ac.uk/Tools/msa/clustalo/). Signal peptides were removed from the alignment.

**Fig. S7.**
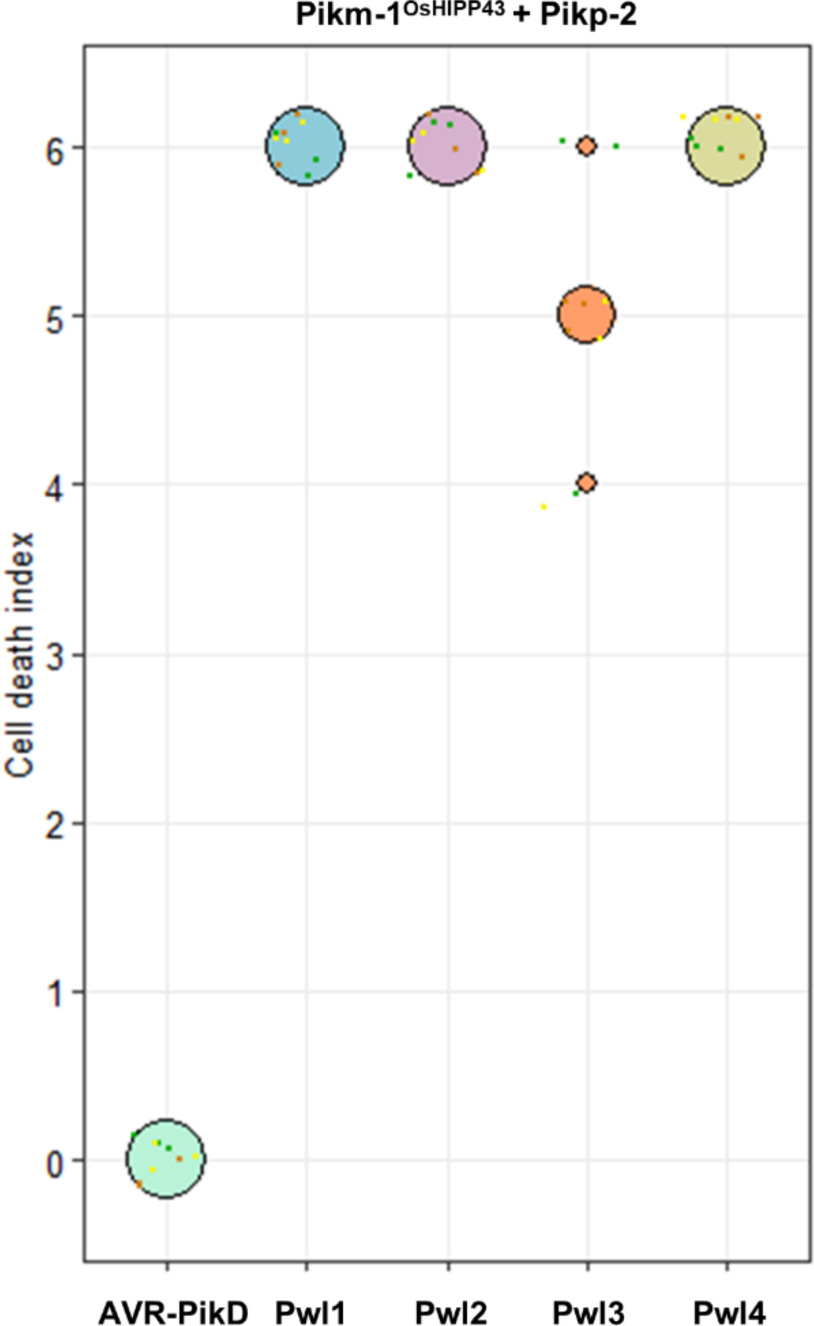
Quantification of cell death assays presented in Fig. 3. Each dot represents a single scoring point in the assay. All dots are randomly scattered around the cell death score for visualisation (size of the circle at given score is proportional to the number of dots within). The colour of each dot reflects independent biological replicates.

**Fig. S8.**
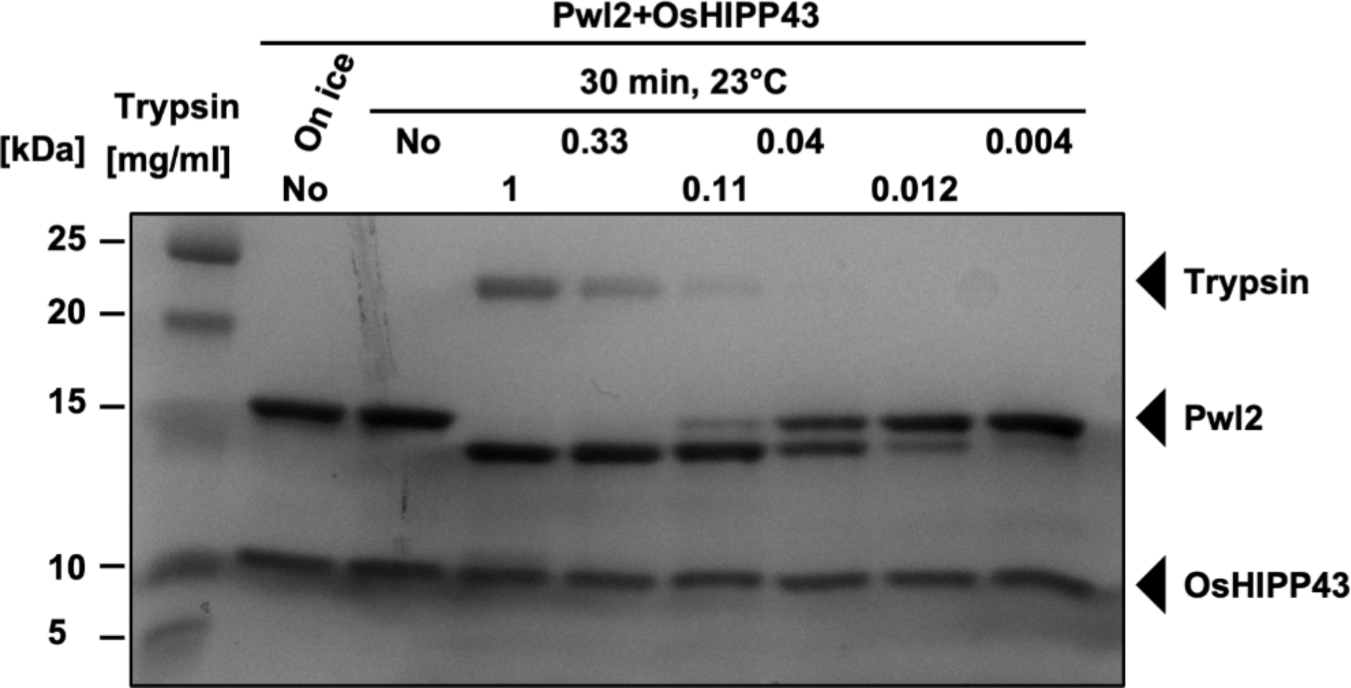
Partial tryptic digest of the purified Pwl2/OsHIPP43 complex. At higher concentrations of trypsin, Pwl2 is cleaved to yield a smaller, stable fragment while OsHIPP43 remains intact.

**Fig. S9.**
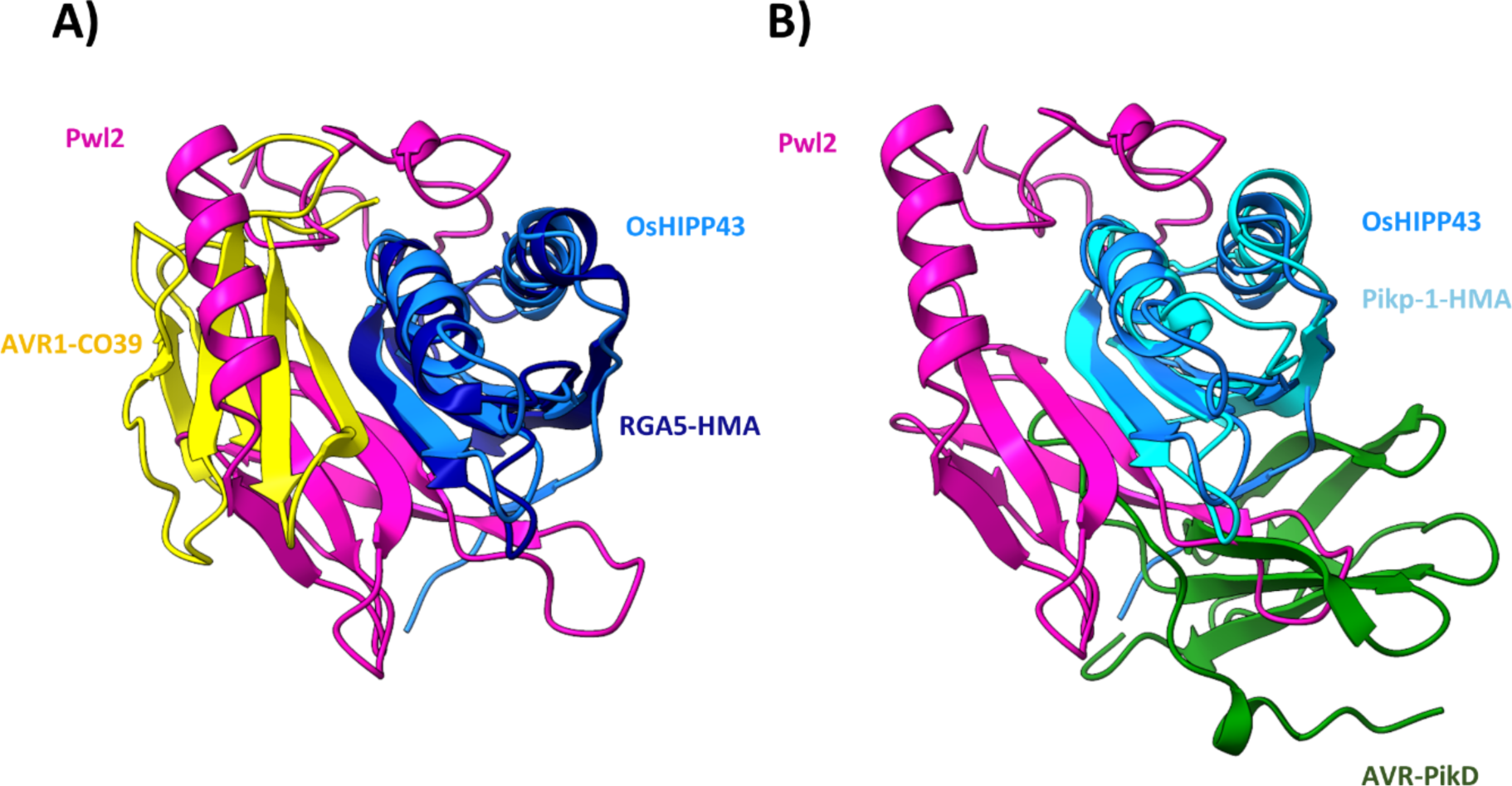
Comparison of the interaction interfaces between Pwl2/OsHIPP43 and selected characterised MAX effector/HMA complexes. A) Pwl2 (pink) / OsHIPP43 (blue) and AVR1-CO39 (yellow) / RGA5-HMA (dark blue) (PDB: 5ZNG). **B)** Pwl2 (pink) / OsHIPP43 (blue) and AVR-PikD (green) / Pikp-HMA (light blue) (PDB: 6G10). The structures were overlaid using ChimeraX (69).

**Fig. S10.**
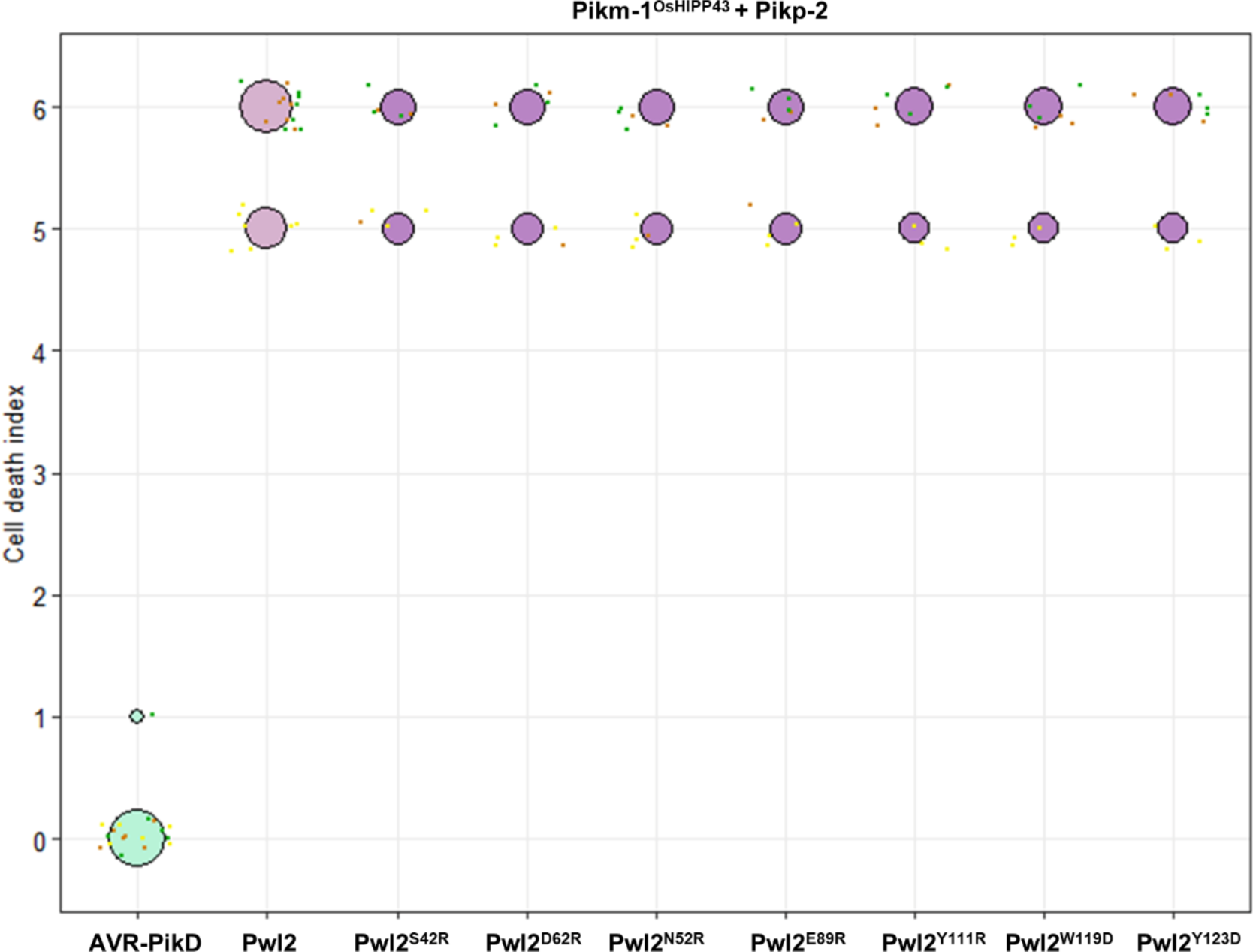
Quantification of the cell death assays presented in Fig. 5A. Each dot represents a single scoring point in the assay. All dots are randomly scattered around the cell death score for visualisation (size of the circle at given score is proportional to the number of dots within). The colour of each dot reflects independent biological replicates.

**Fig. S11.**
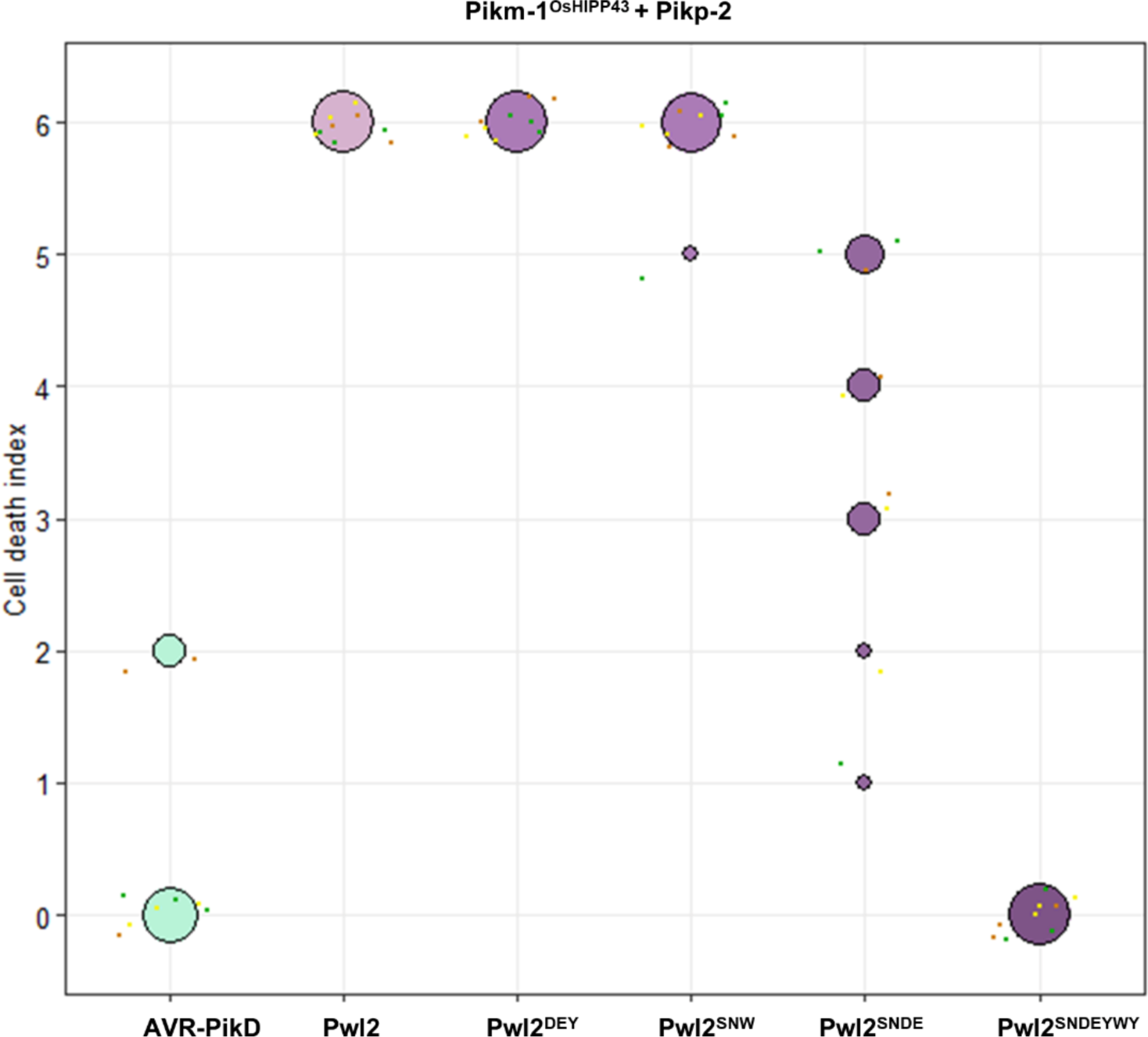
Quantification of the cell death assays presented in Fig. 5B. Each dot represents a single scoring point in the assay. All dots are randomly scattered around the cell death score for visualisation (size of the circle at given score is proportional to the number of dots within). The colour of each dot reflects independent biological replicates.

**Fig. S12.**
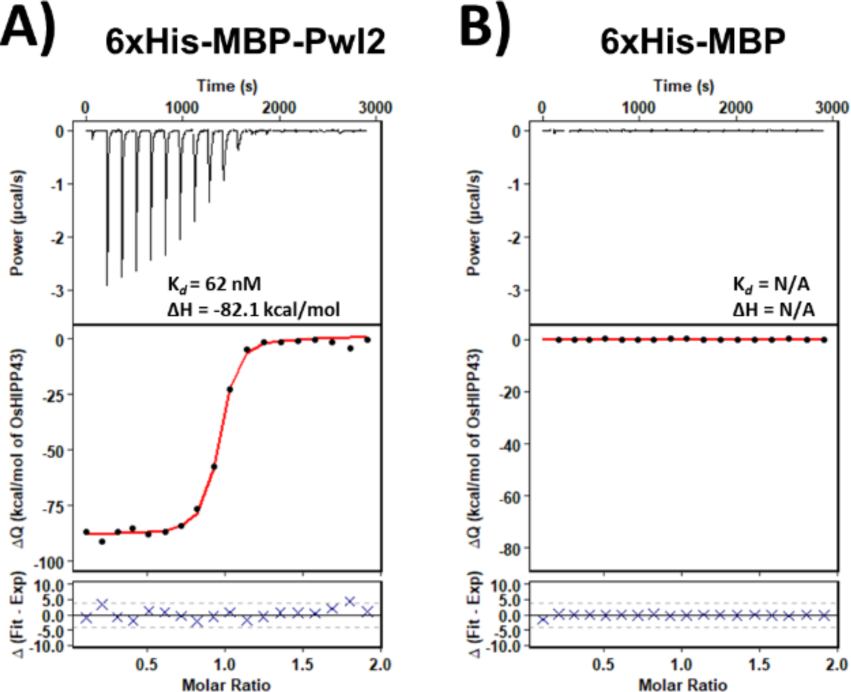
Presence of the 6xHis-MBP-tag does not interfere with the Pwl2/OsHIPP43 interaction measurements recorded using ITC. A) The affinity of 6xHis-MBP-Pwl2 with OsHIPP43 is comparable to Pwl2 alone. **B)** 6xHis-MBP alone does not show affinity to OsHIPP43. **Top panels-** Representative raw isotherm showing heat exchange upon the series of injections of the OsHIPP43 into the cell containing the effector. **Middle panels-** Integrated peaks from the technical replicates and global fit to a single site binding model as calculated using AFFINImeter. **Bottom panels-** Difference between predicted value of measurement (by global fit) and actual measurement as calculated using AFFINImeter.

**Fig. S13.**
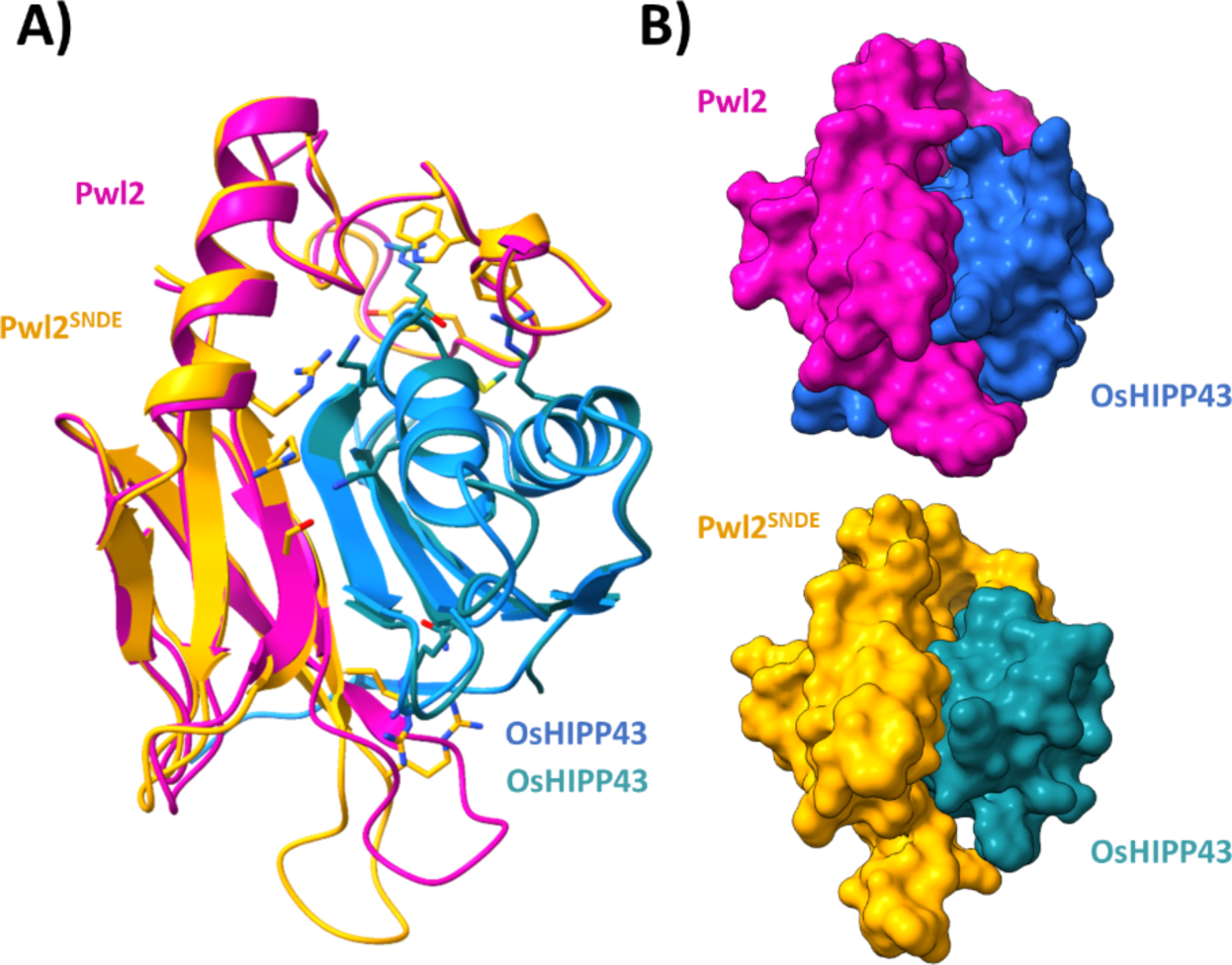
Crystal structure of the Pwl2^SNDE^/OsHIPP43 complex reveals how the mutations are accommodated at the interface. **A)** Superposition of the Pwl2 (pink) / OsHIPP43 (blue) and Pwl2^SNDE^ (orange) / OsHIPP43 (green) complexes, with mutated residues depicted as sticks on the Pwl2^SNDE^/OsHIPP43 structure. **B)** Side-by-side comparison of surface representation of the two complexes reveals how the introduced mutations in Pwl2^SNDE^ affect the OsHIPP43 interaction interface globally. The structures were overlaid using ChimeraX (69).

**Fig. S14.**
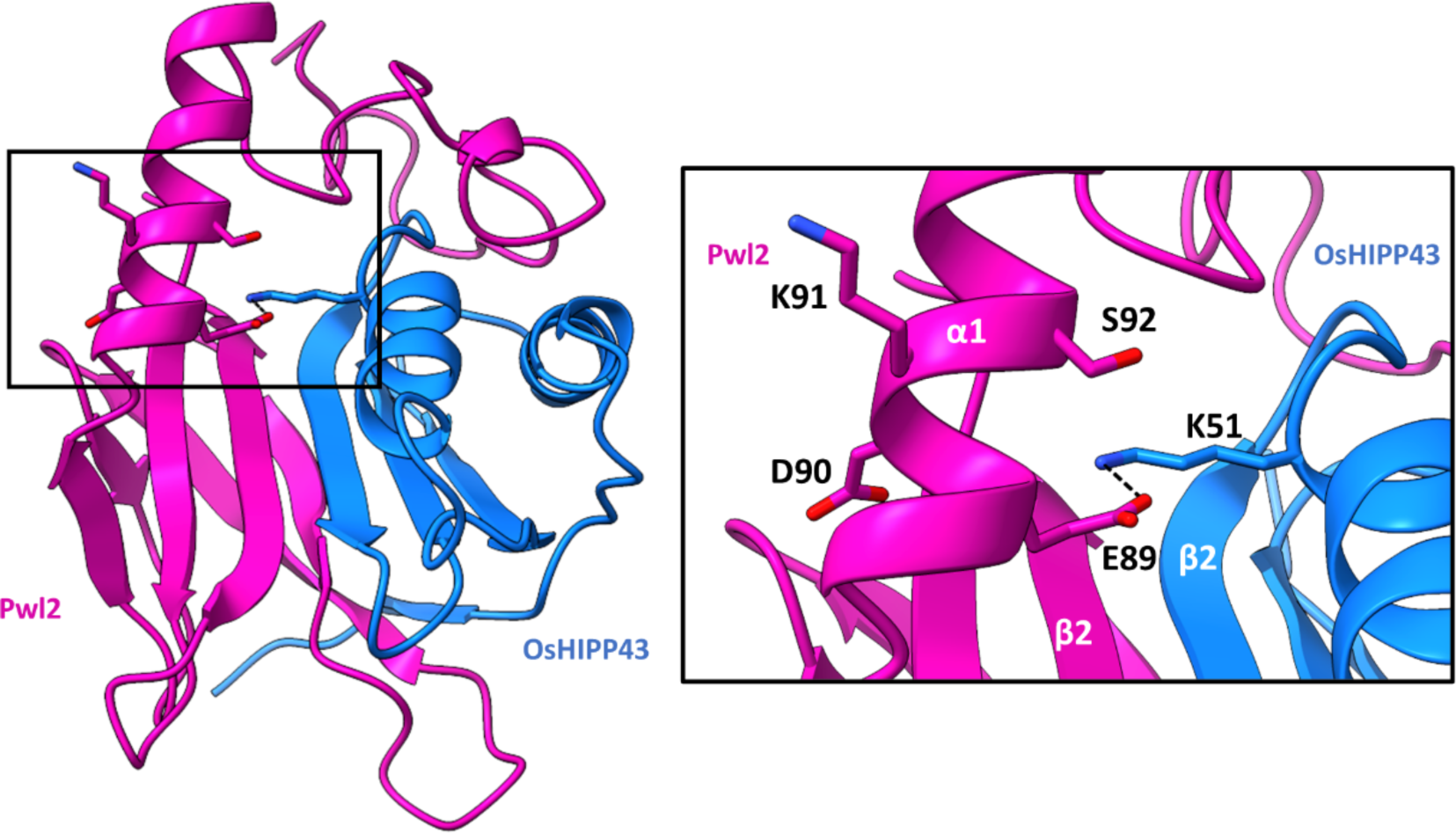
The polymorphic region of Pwl2 is not buried at an interface with OsHIPP43. Residues Glu-89, Asp-90, Lys-91 and Ser-92 are polymorphic between Pwl2 alleles. Close-up view of the polymorphic region within the Pwl2 (pink) / OsHIPP43 (blue) crystal structure shows that only Glu-89 residue is involved in OsHIPP43 (blue) binding. Side chains shown in stick representation. α-helices and β-strands are labelled, and amino acids labelled with single letter codes. Hydrogen bonds are depicted as black dashes between atoms.

**Table S1.** *X*-ray data collection statistics.

|  | PwI2/OsHIPP43 | PwI2/OsHIPP43 | PwI2<sup>SNDE</sup>/OsHIPP43 |
| --- | --- | --- | --- |
|  | Native | S-SAD | Native |
| Wavelength (Å) | 0.9795 | 1.90 | 0.9795 |
| Space group | *P* 6<sub>1</sub> 2 2 | *P* 6<sub>1</sub> 2 2 | *P* 2<sub>1</sub> 2<sub>1</sub> 2<sub>1</sub> |
| Cell dimensions<br>*a*, *b*, *c* (Å) | 63.16 63.16 198.21 | 63.30 63.30 198.47 | 66.38 66.80 81.90 |
| Resolution (Å)* | 54.70-1.80 (1.84-1.80) | 54.82-2.80 (2.95-2.80) | 47.09-2.50 (2.60-2.50) |
| *R*<sub>meas</sub> (%) | 8.7 (169.8) | 13.5 (43.3) | 29.6 (139.7) |
| *I*/σ*I* | 31.1 (2.9) | 70.5 (28.3) | 6.6 (1.9) |
| Completeness (%) |  |  |  |
| Overall | 100.0 (100.0) | 100.0 (99.8) | 100.0 (100.0) |
| Anomalous | 100.0 (100.0) | 100.0 (100.0) | 100.0 (100.0) |
| Unique reflections | 22749 (1291) | 6420 (888) | 13157 (1454) |
| Redundancy |  |  |  |
| Overall | 36.9 (38.4) | 295.3 (230.7) | 13.3 (13.9) |
| Anomalous | 20.3 (20.5) | 173.7 (129.2) | 7.2 (7.3) |
| CC<sup>(1/2)</sup> (%) | 100.0 (91.5) | 99.8 (99.8) | 99.2 (76.5) |

\*The highest resolution shell is shown in parenthesis.

**Table S2.** Refinement and model validation statistics.

|  | PwI2/OsHIPP43 | PwI2<sup>SNDE</sup>/OsHIPP43 |
| --- | --- | --- |
| Resolution (Å) | 54.76-1.8 (1.85 – 1.80) | 47.13-2.50 (2.57-2.50) |
| $R_{\text{work}}/R_{\text{free}}$ (%) | 18.8 (25.1) / 21.5 (29.5) | 21.6 (27.0) / 28.6 (35.0) |
| No. atoms |  |  |
| Protein | 1473 | 2847 |
| Ligand | 175 | 43 |
| B-factors |  |  |
| Protein | 43.64 | 41.88 |
| Ligand | 46.00 | 31.24 |
| R.m.s deviations |  |  |
| Bond lengths (Å) | 0.009 | 0.007 |
| Bond angles (°) | 1.62 | 1.73 |
| Ramachandran plot (%)** |  |  |
| Favoured | 97.3 | 97.2 |
| Allowed | 2.7 | 2.5 |
| Outliers | 0.0 | 0.3 |
| MolProbity Score | 1.3 | 2.0 |

\*The highest resolution shell is shown in parenthesis.
\*\*As calculated by MolProbity

